# Reversal of neuronal tau pathology, metabolic dysfunction, and electrophysiological defects via adiponectin pathway-dependent AMPK activation

**DOI:** 10.1101/2024.02.07.579204

**Authors:** Eric R. McGregor, Danny J. Lasky, Olivia J. Rippentrop, Josef P. Clark, Samantha L. G. Wright, Mathew V. Jones, Rozalyn M. Anderson

## Abstract

Changes in brain mitochondrial metabolism are coincident with functional decline; however, direct links between the two have not been established. Here, we show that mitochondrial targeting via the adiponectin receptor activator AdipoRon (AR) clears neurofibrillary tangles (NFTs) and rescues neuronal tauopathy-associated defects. AR reduced levels of phospho-tau and lowered NFT burden by a mechanism involving the energy-sensing kinase AMPK and the growth-sensing kinase GSK3b. The transcriptional response to AR included broad metabolic and functional pathways. Induction of lysosomal pathways involved activation of LC3 and p62, and restoration of neuronal outgrowth required the stress-responsive kinase JNK. Negative consequences of NFTs on mitochondrial activity, ATP production, and lipid stores were corrected. Defects in electrophysiological measures (e.g., resting potential, resistance, spiking profiles) were also corrected. These findings reveal a network linking mitochondrial function, cellular maintenance processes, and electrical aspects of neuronal function that can be targeted via adiponectin receptor activation.

## Introduction

The specific mechanisms for cognitive decline and dementia associated with Alzheimer’s disease (AD) remain unclear; however, the loss in cognitive performance is more strongly correlated with neurofibrillary tangles (NFT) than with amyloidβ plaques^1,2^. NFTs are composed of aberrant hyper-phosphorylated tau aggregates, a microtubule-associated protein encoded by the MAPT gene. Ordinarily, tau plays a key role in neuronal structure and trafficking of cargo along axons^3,4^. NFTs also accumulate in other neurodegenerative diseases, such as frontotemporal dementia, Parkinson’s disease, and chronic traumatic encephalopathy, so mechanisms underlying NFT formation and interventions for NFT clearance would have broad clinical significance.

Disruptions in metabolic pathways are coincident with the development of AD pathology, whether spontaneous or engineered^5,6^ and tauopathy specifically has been linked to loss of mitochondrial function^7^. Mitochondrial dysfunction is a hallmark of the aging process and may play a causal role in the increase in risk for neurodegenerative disease onset as a function of age. The idea that preserving mitochondrial function might be effective in blunting pathology and improving neural function is gaining traction^8,9^, although precisely how this might be accomplished remains to be shown.

One potential therapeutic target for activating mitochondrial metabolism is adiponectin receptor agonism. Adiponectin is a highly abundant 30 kDa peptide hormone secreted by adipose tissue that regulates metabolism in its target tissues, increasing mitochondrial activity and fatty acid oxidation, and is associated with systemic insulin sensitivity^10,11^. Adiponectin counters metabolic defects linked to obesity^12^, is linked to preserved healthspan in normative aging^13^, and is upregulated by the longevity-promoting intervention of caloric restriction^14,15^. Adiponectin signaling occurs through adiponectin receptors 1 and 2 (adipoR1/2), with a poorly understood contribution from the T cadherin receptor^16^. AdipoRs are expressed in the brain^17^, and recent studies suggest that neuronal adipoR activation may be neuroprotective in the context of diabetes and stroke^18–20^. AdipoRon (AR) is a nonselective adiponectin receptor agonist that binds to both adiponectin receptors 1 and 2 (AdipoR1/2)^21^ and has been shown to promote aggregate clearance in models of AD^22,23^. The present study aimed to investigate the interactions between disease pathology, neuronal function, and neuronal metabolism.

## Results

### AdipoRon clears phosphorylated tau and NFT via AMPK and GSK3b

Primary neurons carrying prion promoter-driven hTauP301S cDNA accumulate phosphorylated Tau and form neurofibrillary tangles, a process that is further augmented by seeding with pre-formed Tau fibrils (PFF). Isolated primary neurons from P0 neonate hTau P301S mouse pups (Fig.S1) were seeded with pre-formed tau fibrils (PFF)^24^ on day seven and cultured for an additional 18–22 days to induce hyperphosphorylated Tau neurofibrillary tangles (NFT) (Fig.1A, upper panels). Treatment with AR (10μM) for 24 hours significantly reduced levels of Tau phosphorylated at S202/T205 and lowered the levels of NFT detected using a tangle-specific antibody (Fig.1A, middle panels)^25^. The best-characterized mediator of adiponectin signaling is AMPK, which is thought to become activated downstream of AdipoR1/2 ligand binding^11^, although most evidence comes from non-neuronal cell types. Treatment with AMPK inhibitor Compound C (8μM) blunted the clearance of hyperphosphorylated Tau in seeded neurons, indicating that reversal of Tau pathology via AR is AMPK-dependent (Fig.1A, lower panels). These data suggest a role for AMPK but do not reveal if it is specifically activated in response to AR in primary neurons. Intracellular signaling by phosphorylation tends to be rapid and transient, prompting an investigation over a shorter timeframe. Seeded primary neurons were treated for 10 minutes with 10μM AR or DMSO control, at which time significantly greater activating phosphorylation at S172 was detected (Fig.1B). Total AMPK was not significantly different in either treated or control neurons over this same time frame (Fig.1B, S2).

**Figure 1:**
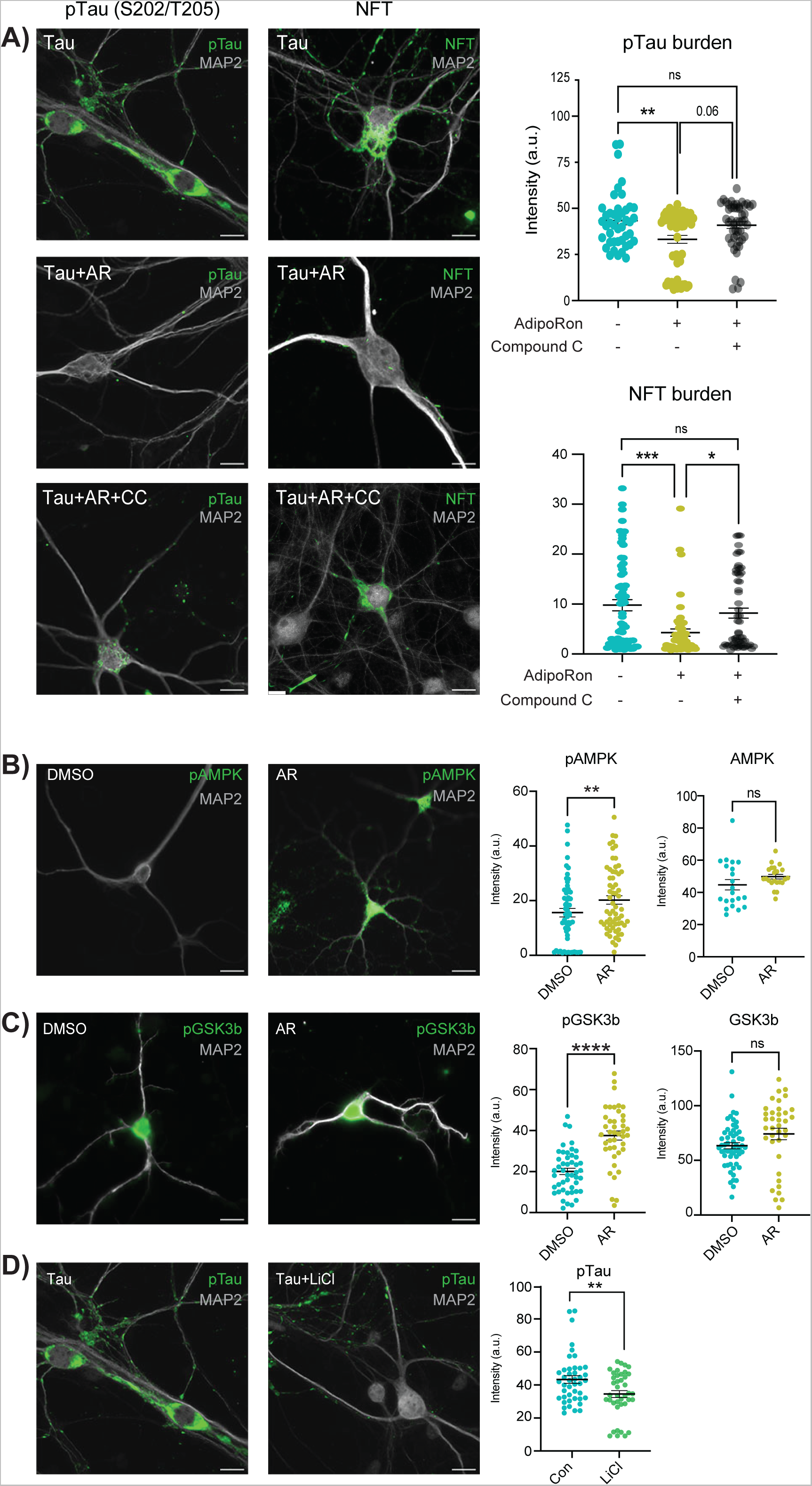
AdipoRon clears phosphorylated tau and NFT via AMPK and GSK3b. A) Immunofluorescence of phosphorylated Tau (pTau) at S202/T205 and NFT (MC-1 antibody) in Tau neurons following 24-hour treatment with 10 μM AdipoRon ± 8 μM AMPK inhibitor Compound C (pTau, n=43–58 cells; NFT, n=47-72 cells). Scale bar 15 μm). B) Quantification of immunofluorescence images in figure A. Data shown as mean ± SEM. Significance determined by one-way ANOVA with Tukey’s multiple comparison’s test. C) Immunofluorescence detection and quantification of phosphorylated AMPK and total AMPK in Tau neurons following 10-minute treatment with 10 μM AdipoRon. (n=52–54 cells, phosphorylated) (n=21-22 cells, total). Significance determined by Student’s t-test with Welch’s correction. C) Immunofluorescence detection and quantification of phosphorylated GSK3b and total GSK3b in Tau neurons following 10-minute treatment with 10 μM AdipoRon. (n=38–55 cells, phosphorylated) (n=36-54 cells, total) Significance determined by Student’s t-test with Welch’s correction. E) Immunofluorescence detection of pTau after 24-hour treatment with 15mM LiCl (n=40-43 cells). Data shown as mean ± SEM. (*:p<0.5, **:p<0.01, ***:p<0.001, ****:p<0.0001).

AMPK is not known to be directly involved in Tau phosphorylation; however, a strong candidate is the established Tau kinase GSK3b (glycogen synthase kinase 3 beta). NFT accumulation has been shown to lead to increased acetylation of GSK3b, modulating its activity and creating a feedforward cycle^26^. Prior independent studies have suggested that it might also play a role in adiponectin signaling^27,28^, and GSK3b is known to be directly inhibited by AMPK^29,30^. Tau neurons treated with AR for 10 minutes contained significantly greater inhibitory phosphorylation of GSK3b at the Ser9 residue (Fig.1C), while total GSK3b remained unchanged (Fig.S2). To confirm prior reports that GSK3β inhibition directly leads to a lower level of phosphorylated Tau, Tau neurons were treated with 15 mM lithium chloride (LiCl), a well-documented GSK3b inhibitor. Within 24 hours of LiCl treatment, phosphorylated Tau (S202/T205) was significantly lower, indicating that GSK3b inhibition is sufficient to reverse Tau modification in primary neurons (Fig.1D). These data place AMPK and GSK3b downstream of AR in clearing the aggregate-prone Tau from pre-seeded primary neurons and show that NFT formation is dynamic and reversible via activation of adiponectin signaling.

### Neuronal transcriptional response to AdipoRon extends beyond metabolic regulation

Initially, AR was developed as a means to correct defects in metabolism linked to obesity ^21^; however, changes in metabolism can have broad cellular effects ^31^. To gain insight into the impact of NFT burden and the response to AR, transcriptional profiling was conducted via RNAseq. RNA was extracted from WT and P301S neurons 18 days post-PFF seeding and following 24-hour treatment with 10μM AR (n=5–6 cultures per group). Almost 1.6 billion sequencing reads were detected (∼78 million reads per sample), trimmed, and aligned to the GRCm39 mouse genome, yielding ∼ 42,000 transcripts from 20,000 unique genes. Differential expression analysis was conducted among the four groups (WT and Tau, with or without AR). Consistent with prior reports, the transcriptional response to the presence of NFTs was negligible^32,33^. Only one gene, Gbp2b (guanylate binding protein 2b), was differentially expressed between Tau and WT neurons and increased 6-fold. Gbp3b is a regulator of inflammatory responses that has been previously associated with neuronal death in response to traumatic injury^34^.

In contrast, transcripts from 138 genes were differentially expressed between the treatment groups (AR vs. control) for Tau neurons (FDR<0.05) (Fig.S3, Table S1). A greater number of significantly differentially expressed genes were detected in AR-treated WT neurons, 656 genes, including 137 of those detected in the Tau neurons for which the directionality of change was conserved (Fig.S3). Top AR-responsive genes were dispersed among cellular functions and included Nptx2 (synaptic communication), Kcnv1 (potassium channel), Sost (Wnt Inhibitor), Ucn (Dopaminergic neuron differentiation), Lyve1 (Membrane glycoprotein), Adcyap1 (Stress response), Pbp2 (MAPK cascade antagonist), Pdcd1 (Immune), and Chrnb3 (cholinergic receptor) (Fig.S3; TableS1). These data show that the impact of AR extends beyond metabolic modulation and suggest that broader aspects of neuronal function were harnessed downstream of adiponectin receptor activation.

Pathway analysis via gene-set enrichment analysis (GSEA) identified AR-responsive pathways in Tau neurons (Fig.2A) and in WT neurons (Fig.2B). Among those shared pathways positively enriched were proteostasis, DNA maintenance and repair, and metabolism. Ribosomal pathways were enriched in both WT and Tau neurons but in opposite directions, perhaps reflecting change in ribosomal composition rather than in ribosomal abundance^35,36^. Homologous recombination and DNA replication pathways were AR-responsive in both genotypes, with base excision repair also detected in the AR-treated Tau neurons. The biology behind this is unclear, and a connection between genome maintenance and adiponectin has not previously been identified.

**Figure 2:**
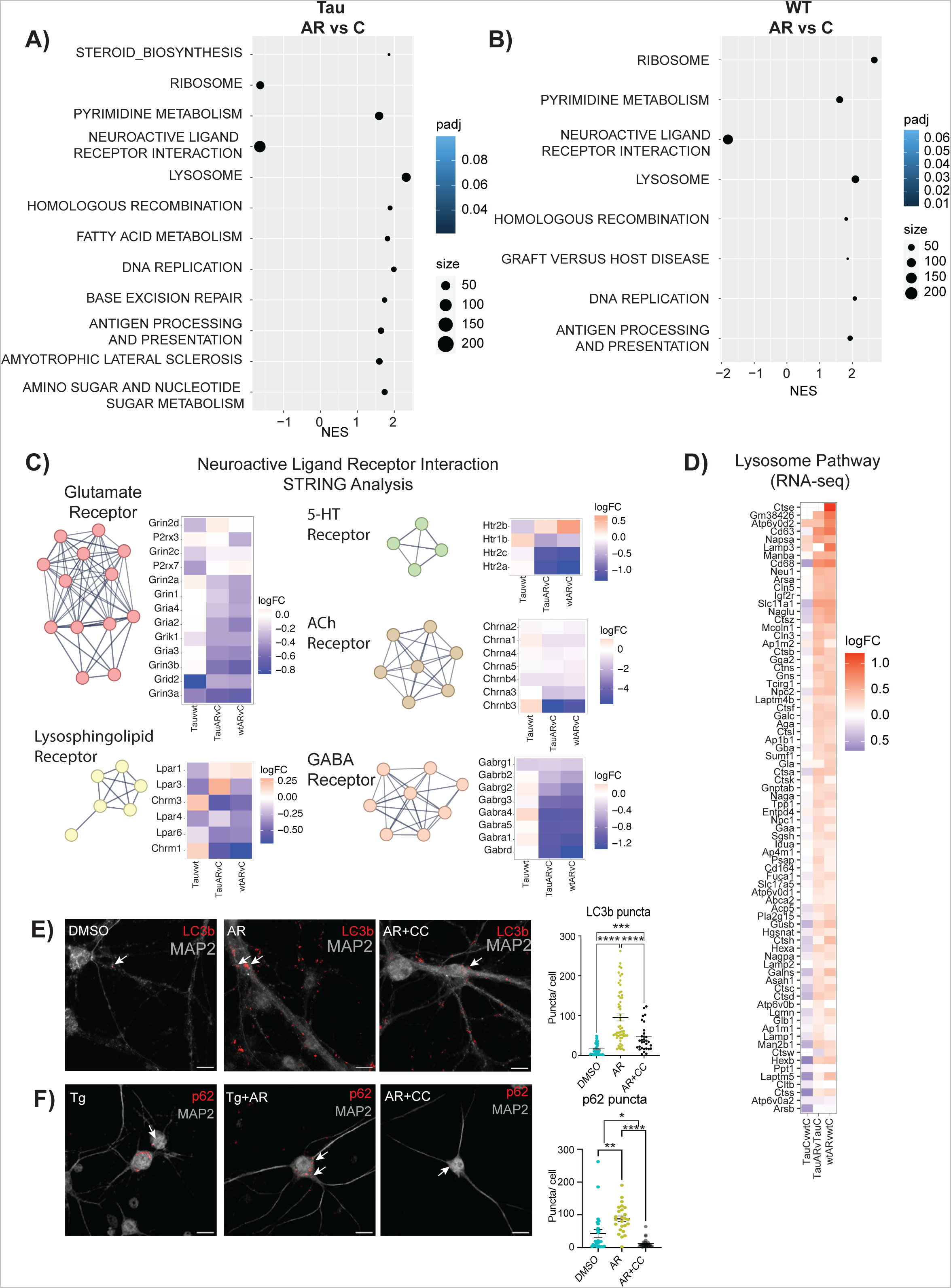
Neuronal transcriptional response to AdipoRon extends beyond metabolic regulation. A) AR responsive KEGG pathways in Tau neurons. Pathway enrichment determined by GSEA. B) AR responsive KEGG pathways in WT neurons. Pathway enrichment determined by GSEA. C) STRING analysis and heatmap displaying log_2_FC of Neuroactive Ligand Receptor Interaction Pathway genes from the Tau neurons in response to AR. Genes were clustered using MCL clusterings. The figure shows all clusters with ≥ 4 genes. D) Heatmap of lysosome pathway detected by GSEA in AR-treated Tau and WT neurons. The log_2_FC of the genes in Tau compared to WT levels was also included. E-F) Immunofluorescent detection and quantification of LC3b and p62 in Tau neurons after 24-hour treatment with 10 μM AdipoRon ± Compound C. Data shown as mean ± SEM. Significance determined by one-way ANOVA with Tukey’s multiple comparison’s test (*:p<0.5, **:p<0.01, ***:p<0.001, ****:p<0.0001) (LC3b,n=33–57 cells p62: n=29-43 cells).

The most highly enriched pathway for both genotypes was neuroactive ligand-receptor interactions. This pathway is composed of the families of receptors involved in synaptic communication as well as many of their ligands. STRING analysis of genes identified within this pathway detected 5 clusters (Fig 2C). The glutamate receptor cluster contained genes encoding six NMDA, three AMPA, one kainate, and one delta subunit. These subunits are associated with ionotropic glutamatergic excitability, suggesting a lowering of excitatory postsynaptic responses with AR, which would protect against excitotoxicity^37^. The GABA_A_ receptor transports Cl^-^ ions, typically inhibiting spike firing by hyperpolarization or shunting. Therefore, a decrease in expression would increase the probability of postsynaptic firing. Nicotinic ACh receptors are also ionotropic, allowing cations to flow and elicit rapid depolarizations. Serotonin (5-hydroxytryptamine, 5-HT) receptors are associated with both metabotropic and ionotropic function and fine-tuning cortical neuron behavior. Serotonin receptor activation has been linked to both amyloid-beta secretion and tau phosphorylation^38,39^. The lysosphingolipid cluster included lysophosphatidic acid receptors (Lpar) and metabotropic acetylcholine receptors. Lpar activation is linked to in vitro neurite retraction, neuronal network formation, and forebrain development^40^, while metabotropic ACh receptor activation leads to the hydrolysis of phosphatidylinositol 4,5 bisphosphate^41^. Together, these transcriptional data suggest that AR impinges on neuronal communication processes, although it is not clear if this is coincident with or a consequence of clearance of the NFT.

### AdipoRon activates the lysosomal pathway via AMPK

Autophagy is a conserved cell process for the degradation of aggregates and dysfunctional organelles. Autophagosome-lysosome fusion allows degradative enzymes of the lysosome to access sequestered protein aggregates and damaged organelles^42^. Autophagy has been proposed as a potential target for alleviating both Aβ plaques and NFTs^43,44^, and lysosomal dysfunction has been implicated in AD progression^45^. Lysosome was one of the most significantly positively enriched pathways in AR-treated Tau and WT neurons (Fig.2D), and there is a hint that this pathway is impacted by Tau, although significance in that case was not reached. AMPK is an established regulator of autophagy^46^, acting through ULK and mTOR to stimulate autophagosome maturation. Activated AMPK increases the formation of autophagosomes that can be visualized by detecting LC3b (microtubule-associated protein 1a/1b light chain 3b) associated puncta^47,48^. Exposure of Tau neurons to AR (10μM) for 24 hours (matching the RNAseq data) significantly increased the number of LC3b puncta (Fig.2E) in an AMPK-dependent manner, with significantly fewer puncta detected in the presence of Compound C (8 μM). The chaperone protein p62 shuttles proteins to the autophagosome for subsequent degradation, and it has been previously linked to the anti-inflammatory effects of adiponectin^49^. A greater number of p62- positive puncta were detected following 24 hours of AR treatment (Fig.2F), again in an AMPK-dependent manner. These data show that autophagosome induction occurs subsequent to AR treatment and suggest that this AMPK-dependent process may contribute to NFT clearance.

### AdipoRon rescues mitochondrial deficits caused by tauopathy

It has been established that metabolic dysfunction occurs in the brain during aging and Alzheimer’s disease^6,50,51^. Effects of AD pathology on mitochondrial function, specifically in the neuron, are less well categorized. PGC-1a is a transcriptional coactivator and master regulator of mitochondrial function that has been indirectly linked to broader cellular processes in diverse cell types, including redox metabolism, cell cycle and cell size determination, arborization, and hypertrophy^31,52–54^. Three promotor regions in the Ppargc-1a gene are active in neurons, including one that is neuron-specific^54^, yielding three transcript variants of PGC-1a. Expression from all three promoters was significantly lower in Tau neurons compared to the WT (Fig.3A).

**Figure 3:**
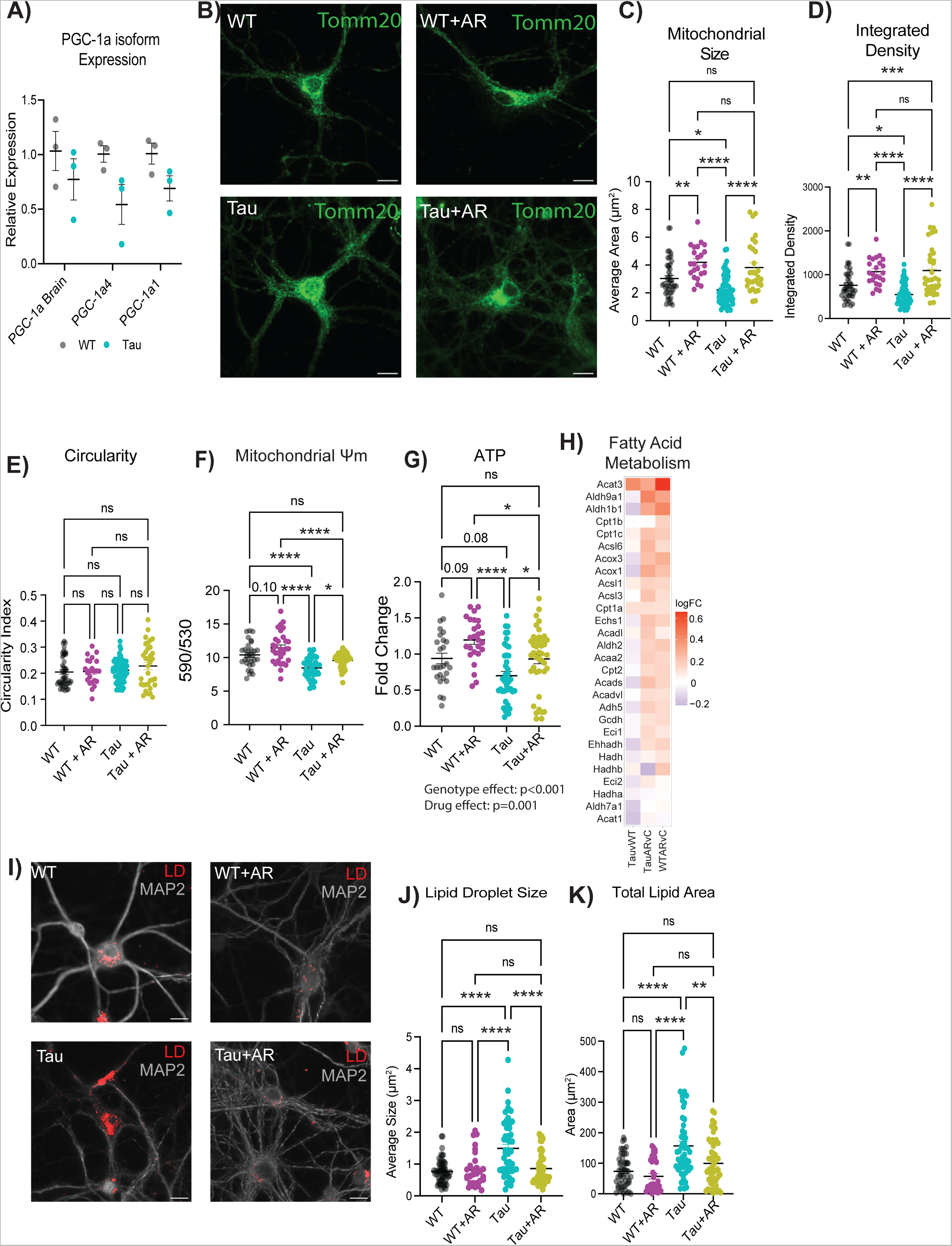
AdipoRon rescues mitochondrial deficits caused by tauopathy. A) mRNA expression of Ppargc-1a isoforms (n=3). B) Representative images of mitochondria detected with TOMM20 antibody. C–E) Mitochondrial morphology was determined through analysis in ImageJ to detect mitochondrial size (C), integrated density (product of mean intensity and mitochondrial area) (D), and circularity (E) (n=25–40 cells). F) Mitochondrial membrane potential measured by JC-1 2 hours after 10 µM AR treatment (n=30-43). G) Relative ATP concentrations after 12 hours of AdipoRon treatment assessed by ATPlite assay (n=26–48). H) Heatmap of fatty acid metabolism GSEA pathway in AR-treated WT and Tau neurons. The log_2_FC of the genes in Tau compared to WT levels was also included for reference. I) Representative images of LipidTox dye (scale bar 15uM). J–K) Quantification of lipid droplet size (J), and total area (K) (n=30–53 cells). Data shown as mean ± SEM. Significance determined by multiple Student’s t-test (A) or two-way ANOVA with Sidak’s multiple comparisons test (C–H, J–L) (* p<0.05, ** p<0.01, *** p<0.001, **** p<0.0001).

Mitochondrial morphology is dynamic and changes in response to metabolic status, with more reticular organization associated with a more oxidative setting^55^. Morphology was assessed by immunofluorescent detection of TOMM20 (translocase of outer mitochondrial membrane 20) (Fig.3B). NFT burden was associated with significantly lower mitochondrial size (Fig.3C) and integrated density (Fig.3D), while the circularity index of the mitochondria was not different (Fig.3E). AR treatment for 24 hours (10μM) restored mitochondrial size and integrated density to that of WT neurons. AR also increased mitochondrial size and integrated density in WT neurons, indicating that the drug may have utility in the absence of AD pathology. Next, the functional consequence of AR treatment on neuronal mitochondrial activity was assessed. Mitochondrial membrane potential is created by electron transport chain activity and drives the ability to derive ATP from oxidative metabolism. Proton motive force quantified by JC-1 assay was lower in Tau neurons (Fig.3F), aligning with the depression in PGC-1a expression. Treatment with AR (10μM) rescued this phenotype within 2 hours and also increased mitochondrial membrane potential in WT neurons. ATP availability was significantly lower in Tau neurons compared to the WT, and AR treatment resulted in significantly greater ATP availability both in Tau neurons and in WT neurons (Fig.3G).

Lipid accumulation is known to be a consequence of inefficient mitochondrial metabolism^29^, and lipid metabolic dysregulation has been linked to neurodegenerative disease progression^56^. AR treatment was associated with a positive enrichment of fatty acid metabolism as detected by RNASeq (Fig.3H). These data suggested that lipid usage and lipid storage could be altered by AR treatment. Lipid droplets (LD) were detected using the neutral-lipid stain HCS LipidTox, revealing differences in lipid stores between WT and Tau neurons (Fig.3I, Fig.S4). Quantitative analysis showed that LD size was significantly greater in neurons with Tau pathology (Fig.3J) and that total lipid stain intensity was significantly greater (Fig.3K). AR treatment lowered LD size and total lipid area, restoring each to WT levels. Together, these data show diminished metabolic capacity and lipid accumulation in neurons with NFT, both of which can be rescued by AR-directed stimulation of the adiponectin pathway.

### AdipoRon restores dendritic complexity through activation of JNK

NFT aggregation is associated with a loss of dendritic spine density and dysfunction in microtubule interactions^57,58^, with implications for neuronal structural integrity. Dendritic arborization can be quantified via immunolabeling for microtubule-associated protein 2 (MAP2), a neuron-specific microtubule protein (Fig.4A). MAP2 was clearly excluded from the nucleus of WT neurons but not Tau neurons. The biological relevance of this difference in subcellular distribution is unclear; however, a similar phenotype has been observed in the brains of Parkinson’s disease patients^59^. Branch detection and scale estimation via Sholl analysis revealed significantly fewer dendritic branches in Tau neurons than in WT, suggesting a loss of dendritic complexity with NFT (Fig.4B). AR treatment (10μM, 24 hours) rescued the NFT-associated compromise in neurite outgrowth, and area under the curve analysis showed significant differences between Tau and WT neurons and between Tau neurons with or without AR. Although the shape of the curve was not identical for WT with or without AR, the change in the distribution of the data was not significant. Soma area was significantly smaller in Tau neurons compared to WT, and AR restored soma size back to WT values (Fig.4C).

**Figure 4:**
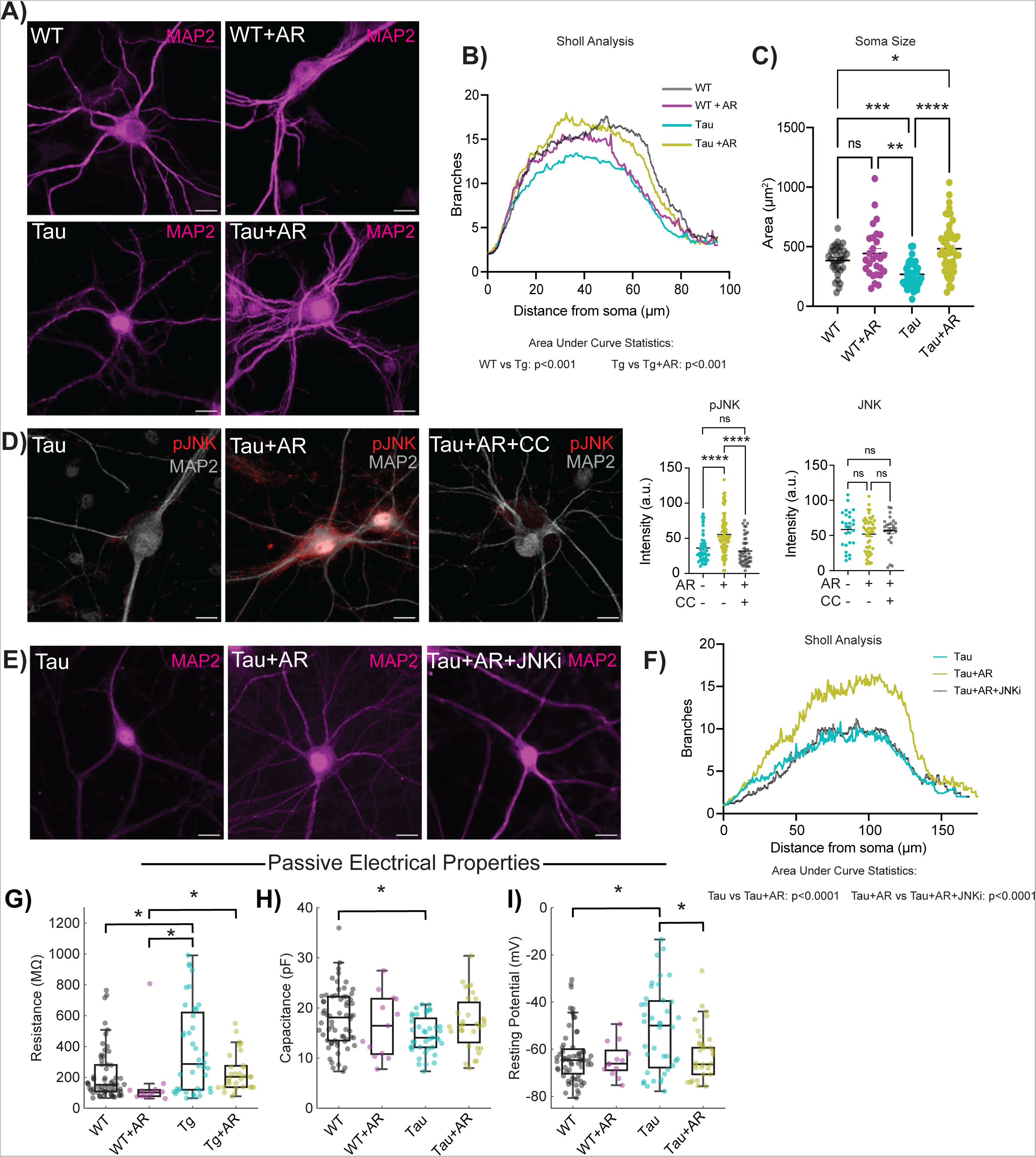
AdipoRon rescues loss of dendritic complexity through activation of JNK. A) Representative immunodetection images of neuronal membranes detected with microtubule-associated protein 2 (MAP2) antibody. Images shown from WT and Tau neurons treated with 10µM AR or DMSO. B) Quantification of neuronal branch number and distance from soma determined by (n = 25–49). C) Quantification of soma size from the same neurons used for Sholl analysis. D) Representative images and quantification of immunodetection of phosphorylated JNK (pJNK) (n=35–51) and total JNK (n=25) in Tau neurons following 10-minute AR treatment (10 µM). E) Representative images of immunodetection of neuronal membranes (MAP2) in Tau neurons treated with 10µM AR ± 10µM SP600125 (JNK inhibitor). F) Quantification of neuronal branch number and distance from soma determined by Sholl analysis (n=47–51). Sholl analysis data shown as mean only. G–I) Passive electrophysiological parameters of intrinsic excitability of neurons. Resistance (G), capacitance (H), and resting potential (I) for WT and Tau neurons with 24-hour AR treatment. Box plots display the median, interquartile range, and whiskers extending to values within the interquartile range multiplied by 1.5. Sample sizes for passive parameters: WT: 64, WT+AR: 13, Tg: 40, Tg+AR: 31. Significance determined by two-way ANOVA with Sidak’s multiple comparisons test (B) or one-way ANOVA with Tukey’s multiple comparisons test (F). Soma size and pJNK/JNK data shown as mean ± SEM. Significance determined by two-way ANOVA with Sidak’s multiple comparisons test or Student’s t-test with Welch’s correction). Significance was determined by Kruskal–Wallis and post-hoc Dunn’s tests. Significance is represented by horizontal brackets between groups (G-I) (* p<0.05, ** p<0.01, *** p<0.001, **** p<0.0001).

The stress-associated kinase JNK (c-Jun N-terminal Kinase) phosphorylates and stabilizes several microtubule-associated proteins, including MAP2 and MAP1B^60^, and has been shown to be required for axonal growth^61^, maintenance of microtubes^60^, and outgrowth of neurons^62^. Adiponectin-induced activation of JNK has been previously reported^63^, but not in neurons, and the mechanism by which this occurs has yet to be established. AR treatment increased activating phosphorylation (T183/Y185) of JNK in Tau neurons (Fig.4D) without impacting overall JNK levels (Fig.S5). Simultaneous treatment with AMPK inhibitor Compound C (8μM) blocked the increase in JNK phosphorylation, placing the phosphorylation and activation of JNK in response to AR downstream of AMPK. Exposure of Tau neurons to JNK inhibitor SP600125 (10μM) for 24 hours blocked the neurite outgrowth induced by AR (Fig.4E). Quantitation of the area under the curve for the distribution of number of branches as a function of distance from soma confirmed that JNK is required for the AR-induced dendritic arborization phenotype (Fig.4F). These data show that JNK is activated by adiponectin receptor pathway in an AMPK dependent manner and is responsible for the AR-driven increase in dendritic complexity.

### Altered passive electrical properties in Tau neurons are restored by AdipoRon

Differences in soma size and dendritic complexity are thought to have functional consequences for the intrinsic electrical properties of neurons^64^. Passive electrical properties of excitable membranes include membrane resistance, capacitance, and resting potential. Whole-cell patch-clamp recordings measure the electrical properties of neurons on a millisecond timescale, providing insight into their excitability. Membrane resistance was significantly elevated in Tau neurons compared to WT, but following 24-hour treatment of 10μM AR, membrane resistance was equivalent to WT neurons (Fig.4G). Capacitance was reduced in Tau neurons compared to WT (Fig.4H), aligning with the reduction in cell body size and dendritic branching, and was also restored to WT by AR. The resting potential is influenced by resting conductances and transmembrane ionic gradients. Tau neurons were relatively depolarized compared to WT (Fig.4I), but AR treatment restored Tau neuron resting potential to WT values (Fig.4I). Collectively, these results suggest that Tau neurons have altered passive electrical properties and that these impairments are restored by treatment with AR. The impact of NFT on electrical properties in neurons raised the possibility that this phenotype might extend to other aspects of AD pathology. Amyloid beta accumulates in the brain and vasculature with AD ^65^, and this pathology can be induced in transgenic mice that express a chimeric mouse/human amyloid precursor protein carrying the 695 Swedish mutation (APP^SWE695^) and a mutant human presenilin 1 (PS1-dE9). There were no statistical differences in membrane resistance, capacitance, or resting potential between APP/PS1 and WT primary neurons (Fig.S5), arguing that electrophysiological defects in primary neurons are specific to NFT.

### Active electrical properties are altered in Tau neurons and restored by AdipoRon

Active electrical properties involve voltage-gated channel functions that underlie action potentials. In patch-clamped neurons, action potentials (Fig.5A-C) elicited in response to stepwise current injections inform of intrinsic excitability. The spiking response at rheobase (Fig.5B), defined as the minimum current application necessary to elicit an action potential, can be used to compute spike-related parameters (Fig.5C). Differences in spike traces following sequential current application (30 steps from least to most depolarizing) were detected between Tau and WT, (−100, −20, 160, 320, 480pA shown), and in response to AR for both genotypes (Fig.5A). There was no statistical difference between WT and APP/PS1 neurons in the latency of their first spike at rheobase, spike amplitude, or width (Fig.S6).

**Figure 5.**
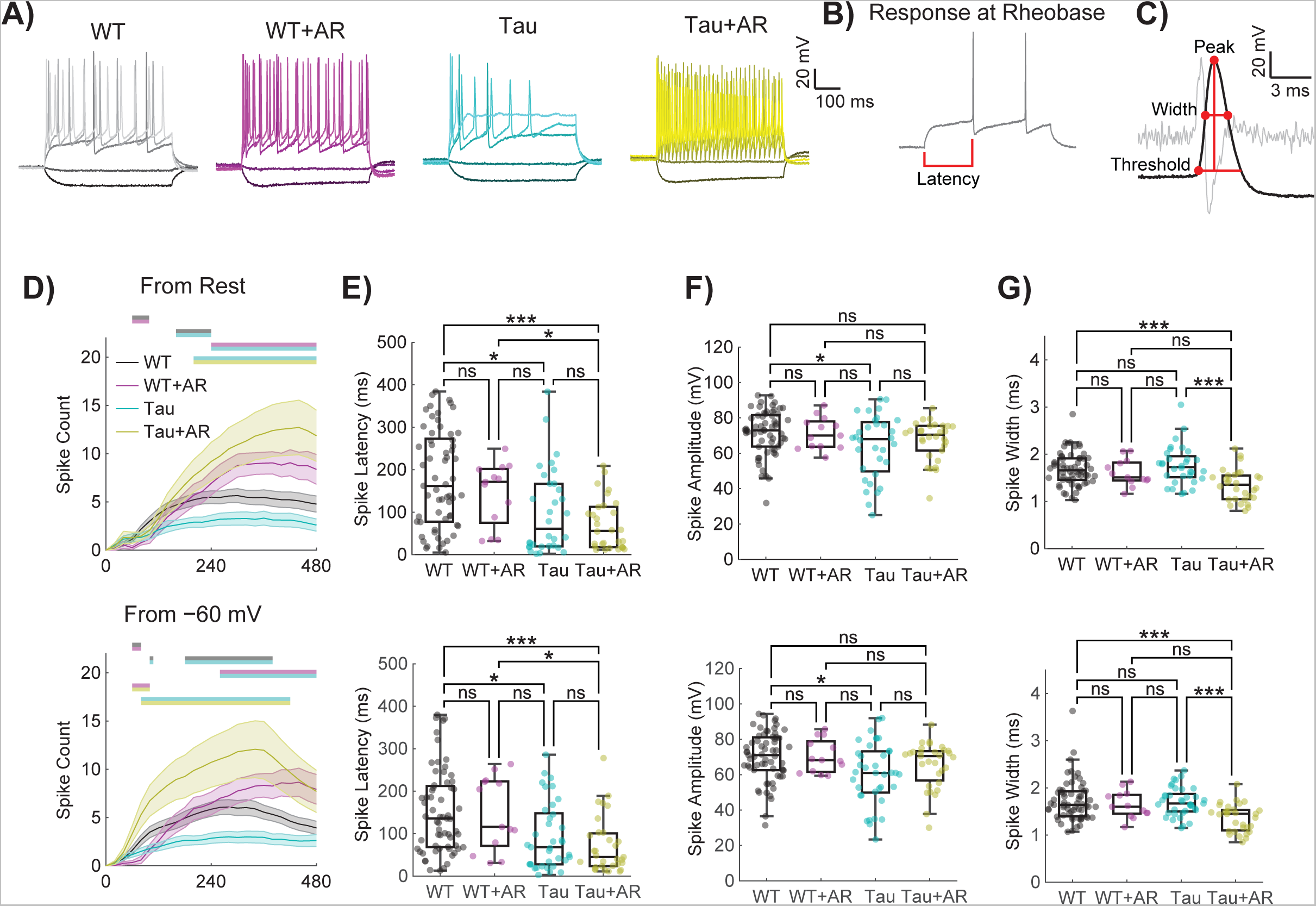
Active electrical properties are altered in Tau neurons and restored by AdipoRon. A) Representative spiking patterns for WT and Tau neurons following 24-hour 10µM AR treatment in response to current applications of −100, −20, 160, 320, and 480 mV (darker to lighter shades). B) Voltage response at rheobase (minimum current required to fire an action potential) for a WT neuron. C) A zoomed version of the first spike fired at rheobase displayed in B. The second derivative is displayed in light gray. The max in the second derivative within a specified window (from when the first derivative was greater than 30 mV/ms to 3 ms prior) determined the threshold voltage for a spike to be fired. Computed rheobase parameters are displayed by the red dots and lines. D) Action potential spike counts were measured in response to 30 stepwise current applications of 20 pA from −100–480 pA for each cell from rest (top) and from −60 mV (bottom). Statistics were performed using repeated measures two-way ANOVA with Geisser–Greenhouse correction and post-hoc Tukey tests. Locations of significantly different comparisons are color-coded by bars at the top. Error bars are SEM. E–G) Spike latency (E), amplitude (F), and width (G) for the first spike fired at rheobase were similarly collected from current-clamp protocols performed for each cell from rest (left) and from −60 mV (right). Spike width was measured using the full width at half amplitude. Box plots display the median, interquartile range, and whiskers extending to values within the interquartile range multiplied by 1.5. Statistics were performed using Kruskal–Wallis and post-hoc Dunn’s tests. Significance is represented by horizontal bars between groups. Sample sizes for D–F are WT: 63, WT+AR: 13, Tg: 37, Tg+AR: 31. (* p<0.05, ** p<0.01, *** p<0.001, **** p<0.0001).

Differences among WT and Tau neurons with respect to AR treatment were further investigated across stepwise current applications in 20 pA increments from −100 to 480 pA (Fig.5D). A significant effect of genotype, treatment, and current step was detected in spike count as measured starting from the resting potential (Fig.5D-G, top row). Since Tau neurons rested relatively depolarized compared to WT (Fig.4I), a “standardized” spike count was also measured by holding near −60 mV (Fig.5D-G, bottom row). Tukey multiple comparisons revealed that Tau neurons had reduced spike counts compared to all other groups, both from resting potential and −60 mV (Fig.5D). Notably, Tau neurons sitting at their native resting potential were less likely to fire one or more spikes at higher current step applications, and from their resting potential and from −60 mV, tended to fire only a single spike unlike the WT or AR treated neurons (Fig.S5). AR-treated neurons of both genotypes were less likely to fire only a single action potential than untreated WT neurons for both recording conditions (Fig.S6).

The neuronal intrinsic excitability and spike morphology can be assessed by analyzing action potentials elicited at rheobase. At both resting potential and from a holding potential of −60 mV, Tau neurons firing action potentials at rheobase were quicker to fire a spike from the onset of the current application (Fig.5E). A short spike latency persisted for Tau neurons after AR treatment. Furthermore, Tau neurons had a reduced spike amplitude at the rheobase compared to WT neurons (Fig.5F). Following AR treatment, Tau neurons fired spikes that were no longer significantly different in amplitude from WT neurons (Fig.5F). Finally, Tau neurons with AR treatment had a reduced spike width compared to WT or Tau neurons without AR treatment (Fig.5G). There were no statistical differences in the spike count across current steps from rest between WT and APP/PS1 neurons (Fig.S6). The results were similar for spike count across current steps from holding at −60 mV, nor were there differences at native resting potential in their tendency to fire one or more spikes across current steps (Fig.S6). These data reveal pervasive disruption in the electrical properties of neurons specific to NFT and provide evidence that at least some of the functional defects associated with Tau pathology can be corrected by treatment with AR.

### AdipoRon stimulates the activity of voltage-gated channels in Tau neurons

The differences in intrinsic electrical properties associated with Tau NFT and their rescue by AdipoRon prompted an investigation of voltage-gated ion channel activity. The activity of such channels can be quantified by controlling the neuronal membrane potential (i.e., voltage-clamp) and measuring the currents induced at different voltages and times. Although we are aware that highly arborized neurons cannot be accurately voltage-clamped with current methods, we nevertheless found informative and statistically significant differences between genotypes and treatments (Fig.6).

**Figure 6.**
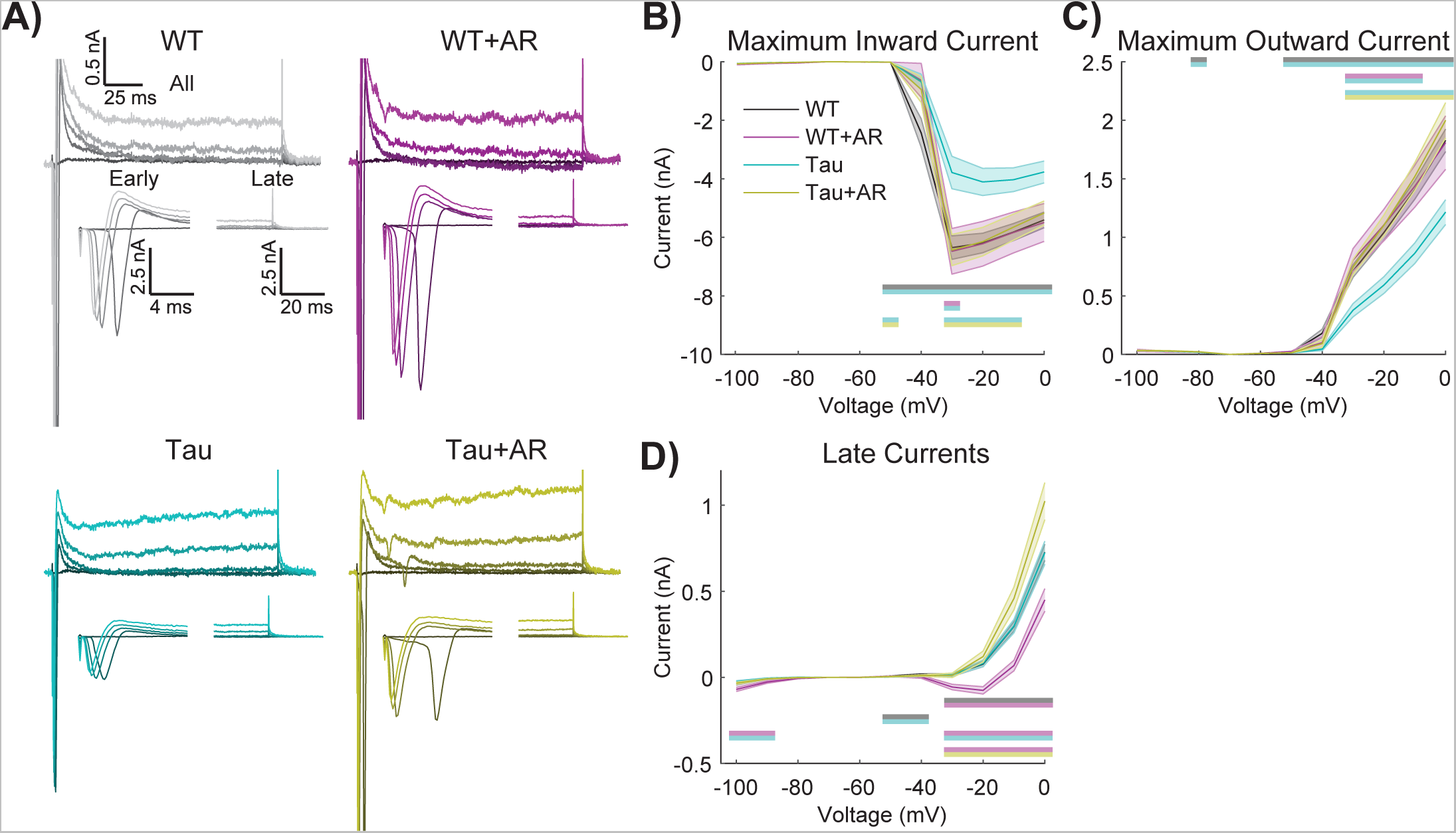
AdipoRon stimulates the activity of voltage-gated channels in Tau neurons. A) Representative current responses of Tau and WT neurons following 24-treatment with 10µM AR (or DMSO) in response to voltages of −40, −30, −20, −10, and 0 mV (darker to lighter shades). B–D) The maximum inward current (B), maximum outward current (C), and IV curves (D) in response to stepwise voltage injections of 10 mV from 0–100 mV. Statistics were performed using repeated measures two-way ANOVA with Geisser–Greenhouse correction and post-hoc Tukey tests. Significant differences are indicated by color-coded by horizontal bars. WT and Tau IV curves are almost visually indistinguishable. Sample sizes are WT: 64, WT+AR: 13, Tg: 40, Tg+AR: 31.

Using a range of applied voltages from a holding potential of −60 mV (Fig.6A), the maximum inward current (i.e., likely an unclamped Na+ current) was measured for untreated and AR-treated WT and Tau neurons (Fig.6B). Tukey multiple comparisons revealed that Tau neurons had a reduced maximum inward current compared to all other groups. Significant main effects of genotype and treatment were detected, where AR had no effect on WT neurons but restored activity in Tau neurons to that of WT (Fig.6B, Table S2). Maximum outward current (i.e., probably a fast, transient potassium current, Fig.6C) of Tau neurons was less than all other groups, with significant main effects of genotype and treatment. AR had no effect on maximum outward current in WT neurons but restored activity in Tau neurons to that of WT.

The maximum inward and outward currents occur within the first few milliseconds (i.e., “Early” in Fig.6A). However, smaller, slower yet persistent currents (i.e.,. “Late” in Fig.6A) are also likely to be important. These persistent currents were not different between WT and Tau neurons; however, the response to AR was genotype-specific (Fig.6D). Multiple comparisons testing revealed that WT neurons with AR treatment had significantly larger net inward currents compared to all other groups (Fig.6D). Voltage-clamp protocols showed no significant differences between WT and APP/PS1 neurons (Fig.S7). Together, the differences in inward/outward current ratio suggest that voltage-gated channel activity is compromised in Tau neurons and that AR treatment tends to favor greater intrinsic excitability.

## Discussion

Data presented here show the neuroprotective and restorative effects of the nonselective adiponectin receptor agonist AR in NFT-burdened primary cortical neurons. Mechanistically, this effect was shown to be AMPK-dependent and linked to the actions of Tau kinase GSK3b. Additional mechanisms for NFT clearance induced by AR include lysosomal genes, an observation that was corroborated in vitro by increases in puncta-associated p62 cargo protein and LC3b, a major component of mature autophagosome formation. Changes in autophagosomes were also dependent on AMPK. Energetic defects linked to NFT were restored by AR, including mitochondrial membrane potential and ATP availability. Neuronal functions such as synaptic communication, axonal transport, vesicle synthesis, dynamic arborization, and maintenance of electrophysiological capacity create energetic and anabolic demands. It is unclear if the reduced availability of ATP is a major driver in NFT-associated loss of neuronal function or if other metabolic outcomes also play a role. For example, this study shows that Tau neurons display aberrant lipid accumulation. The greater lipid burden may be linked to inefficient or ineffective lipid fuel use. The positive enrichment of fatty acid metabolism genes with AR treatment is consistent with this idea. Another possibility is that blunted lipid mobilization impacts membrane-related cellular functions. Prior studies have linked lipid remodeling to adiponectin receptor signaling in kidney cells^66^, raising the possibility that the consequences of lipid accumulation in Tau neurons are not limited to problems of energy derivation.

This study shows reduced dendritic complexity in Tau neurons. The rescue of this phenotype by AR was JNK-dependent and AMPK-dependent. Although not tested in this study, it will be interesting to see if changes in arborization impact inter-cellular communications. Given the surprising lack of differences between WT and Tau neurons at the gene transcription level, it is unclear if synaptic processes, including assembly and presentation, are compromised by NFT. AR downregulated genes involved in synaptic transmission, an adjustment that, together with the increase in dendritic complexity, would be predicted to influence cell-cell communication. In primary hippocampal neurons, arborization and synaptic density are reliant on sufficiency in ATP production^67^. Electrophysiological properties are also heavily influenced by ATP availability. Resting potential depends mainly on the potassium gradient, set up by the ATP-dependent Na^+^/K^+^-ATPase, and on the resting K+ conductance, set up by the number and function of K+ channels. Voltage clamp experiments revealed that Tau neurons had reduced maximum inward and outward currents, but following 24-hour AR treatment, they had maximum inward and outward currents that matched WT. Differences in voltage-gated channel expression were not detected at the mRNA level, indicating that functional differences could arise from changes at the protein or posttranslational level^68^. Impaired K+ channel function would contribute to higher input resistance and less healthy resting potential, whereas impaired Na+ channel function would contribute to defects in active properties (spiking), all of which were observed for Tau neurons. Tau neurons were less capable of generating action potentials and had altered spike properties. AR moved several of these spike properties to align with WT neurons. Ion channel production, localization, and function are all energy-dependent processes, indicating that rescue may be occurring via the improvement of cell metabolic health and ATP availability. The neuroactive ligand-receptor interaction pathway was the most enriched AR-responsive pathway and was negatively regulated in both WT and Tau neurons. Adiponectin is largely thought to modulate cellular metabolism, so this finding was unexpected. The functional significance of this adaptation is not clear but may be linked to changes in action potential firing and the avoidance of excitotoxicity caused by increased neurotransmitter release.

In contrast to the Tau neurons, we observed little impact of APP/PS1 in neuronal passive or active intrinsic excitability; however, this lack of difference in intrinsic electrical properties does not preclude synaptic communication along neuronal networks being disrupted by extracellular plaques. For Tau neurons, the electrophysiological defects reported here are coincident with defects in metabolism, proteostasis, soma size, and dendritic architecture. The intersectionality of these cellular phenotypes is not known. The AR-directed clearance of NFT is associated with mitochondrial rescue, increased available ATP, and activation of proteostasis. One possibility is that AR activates a regulatory hub that encompasses each of these phenotypes. Along these lines, studies in non-neuronal cells show that small changes in mitochondrial function have a broad impact on cellular processes related to fuel preference, energy storage, growth, and homeostasis^31^. Perhaps changes in mitochondrial function that occur within 2 hours are the trigger to induce changes in other cellular processes, or maybe other aspects of intracellular signaling induced by AR create the framework for a coordinated response.

Although these findings are promising, there are important caveats. First, our data are limited to primary cell culture. In reality, neurons operate in an environment of diverse cell types, each with the ability to support and influence neuronal function. It will be important to extend the work to in vivo models, where neuronal function is assessed in situ. Two recent papers indicate that AR is also effective in ameliorating the impact of AD pathology in vivo^23,69^, but the rescue of neurofunctional abnormalities and neuronal excitability defects^70^ in vivo remains to be established. The second limitation is that our study did not include an analysis of Tau isoform expression. Alternate splicing produces different isoforms in humans that differ structurally and regionally in their accumulation in a disease-specific manner^71^, but none of that biology is interrogated in this study. Another limitation is that the use of the transgenic seeded neurons reflects the condition of tauopathy but cannot inform about the etiology of the condition. Nonetheless, the treatment of primary Tau neurons with AR, having been inspired by studies of metabolic regulation in the context of caloric restriction, has demonstrated a broader beneficial effect than had been anticipated. Taken together, our data show that NFT impinges on multiple aspects of neuronal health and function and suggests that AR, through its actions on mitochondrial metabolism, autophagy, and neuronal excitability, is a promising therapeutic for AD and related tauopathies.

## Acknowledgments

This work was supported by NIH AG057408, AG067330, T32DK007665 (ERM), and the Simons Foundation. This study was conducted using resources and facilities at the William S. Middleton Memorial Veterans Hospital, Madison, WI, and the University of Wisconsin-Madison Department of Neuroscience.

The authors declare no conflict of interest.

## Methods

### Cell culture

#### Primary Neurons

Primary cortical neurons were isolated from brains of P0 hTau P301S and APP/PS1 mice. Briefly, the cortices were harvested from pups into ice-cold HBSS where the midbrain and meninges were removed. Brain tissue was minced and digested in 0.25% trypsin for 20 minutes at 37 °C. Trypsin was quenched with DMEM/10%FBS/1%Pen/Strep and cells were dissociated and counted prior to plating on poly-d-lysine coated plates. Cultures were then fully changed to Neurobasal Plus Media (2% B27^+^, 1% GlutaMax, 1% Pen/Strep) the subsequent day. Upon first change to Neurobasal media, neurons were treated with 1 μM AraC (cytosine arabinoside) to eliminate glial cell populations. Media was changed by ½ volume every 3–4 days until experiments. To induce neurofibrillary tau tangle accumulation in the P301S, transgenic neurons were seeded with 3 μg/mL of pre-formed fibrils (PFF) (StressMarq) on day 7 *in vitro* and then cultured for 18 days before analysis ^27,28^ All primary neuron experiments were conducted between day 25–30 *in vitro*. APP/PS1 neuron cultures showed elevated levels of Aβ and, therefore, were not seeded with exogenous protein.

#### Drug Treatments

Cells were treated with 10 μM AdipoRon or DMSO (vehicle control) prior to all experiments as shown above. Treatment was accompanied by a 50% media change. For 10-minute treatment, AdipoRon was added as a 5X solution diluted to a final 1X in the culture media. Compound C (Millipore Sigma; 171260) treatment was co-administered with AR during 50% media change as an 8 μM dose. SP600125 (MedChem Express; HY-12041) was co-administered with AR during 50% media change as a 10 μM dose. Lithium chloride (Sigma Aldrich; L7026) was administered for 24 hours through a 50% media change at a final concentration of 15 mM.

### Immunofluorescence

Conducted using standard techniques. Immunofluorescence images were acquired on a Leica DMi8 inverted microscope equipped with a HCX PL Fluotar 100X /1.30 oil objective. Images were analyzed using ImageJ (NIH, Wayne Rasband, http://rsb.info.nih.gov/ij/). Cells were grown on glass coverslips coated with poly-d-lysine. Cells were fixed in 10% formalin for 10 minutes, permeabilized with 0.3% Triton X-100 in PBS for 1 hour, and incubated with primary antibody cocktail overnight. Primary antibodies used: MAP2 (ab5392; Abcam), AT-8 (MN1020; Thermo Scientific), MC-1 (Peter Davies lab), LC3b (ab48394; Abcam), p62 (23214; Cell Signaling Technologies), pGSK3b (Ser9) (9336; Cell Signaling Technologies), GSK3b (9832; Cell Signaling Technologies, pAMPK (Thr172) (2535; Cell Signaling Technologies), AMPK (2532; Cell Signaling Technolgies), Tomm20 (ab56783; Abcam), pJNK (Thr183/ Tyr185 (9255; Cell Signaling Technologies), and JNK (9252; Cell Signaling Technologies). Lipid droplets were detected using HCS LipidTox Red following manufacturer’s protocol (H34476; Thermo Scientific).

Sholl analysis was conducted using the SNT plugin (https://imagej.net/plugins/snt/) for ImageJ.

### RNA sequencing

RNA extraction was completed using a Direct-zol RNA kit (Zymo Research, Irvine, CA) according to the manufacturer’s instructions. Each RNA library was generated following the Illumina TruSeq RNA Sample Preparation Guide and the Illumina TruSeq RNA Sample Preparation. Purified total RNA was used to generate mRNA libraries using NEBNext Poly(A) mRNA Magnetic Isolation Module and NEBNext Ultra RNA Library Prep kit for Illumina (Illumina Inc., San Diego, CA, USA). Quality and quantity were assessed using an Agilent DNA1000 series chip assay and Invitrogen Qubit HS Kit (Invitrogen), respectively. Sequencing reads were trimmed to remove sequencing adaptors and low-quality bases (Jiang et al., 2014), aligned to mm10 reference genome using the STAR aligner, and alignments used as input to RSEM for quantification. Differential gene expression analysis was performed via EdgeR generalized linear model (GLM) method.

#### String analysis

MCL clustering was used with an inflation parameter of 3. Active interaction sources used: Experiments, databases, co-expression, neighborhood, gene fusion, co-occurrence. The minimum required interaction score was 0.700.

### ATP luminescence assay

Relative ATP levels were quantified with the ATPlite assay (Perkin-Elmer, Waltham, MA). Luminescence was measured using an Infinite M200 microplate reader (TECAN, Grodig, Austria). Neurons were cultured as above and treated with vehicle or AdipoRon (10 μM) for 12 hours.

### JC-1 Assay

Mitochondrial membrane potential was quantified with JC-1 dye (Invitrogen, Waltham, MA). Neurons were treated for 2 hours and then incubated with 1ug/mL JC-1 dye for 30 minutes. Cells were washed with PBS prior to fluorescence detection. Fluorescence was measured using excitation/emission wavelengths of 535/590 nm and 485/530 nm.

### qRT-PCR

Cells were treated as outlined above and RNA was collected 24 hours after treatment. For NSC differentiation and primary neuron maturation, cells were lysed at day in vitro indicated above. Cells were lysed with Trizol and RNA was isolated using Zymo Research Direct-zol RNA MiniPrep Kit. RT-qPCR was conducted using iTaq Universal SYBR Green Supermix (1725121, Bio-Rad, Hercules, CA, 94547). Primer sequences for all transcripts can be found in Table S2.

### In Vitro Electrophysiology Recordings

All recordings were done in extracellular solution containing (in mM): 145 NaCl, 2.5 KCl, 1 MgCl_2_, 2 CaCl_2_, 10 HEPES, and 10 dextrose. Extracellular solution was adjusted to a pH of 7.3 with 5N NaOH and 320–325 mOsm with sucrose. Whole-cell patch-clamp recordings were made using an upright microscope (Axioskop FS2, Zeiss) with infrared differential interference contrast optics. Patch pipettes pulled from thin-walled borosilicate glass (World Precision Instruments) had a resistance of 3–5 MΩ when filled with intracellular solution containing (in mM): 135 K-gluconate, 5 KCl, 0.1 EGTA, 10 HEPES, 2 MgATP, 0.3 Na_2_GTP, 0.25 CaCl_2_, and 20 Na_2_-phosophocreatine. Intracellular solution was adjusted to a pH of 7.2 with 5 N KOH and 310–315 mOsm with H_2_O.

Recordings were done using an Axopatch 200B or MultiClamp 700B amplifier (Axon Instruments), filtered at 5 kHz using a 4-pole Bessel filter, and digitized at 10 kHz using a Digidata 1320A or 1322A analog-digital interface (Axon Instruments). Data were acquired to a Power Mac G4 (Apple) using Axograph X v1.5.4 (Axograph.com). Input resistance, series resistance and capacitance were assessed from the fit of the electrical responses to −5 mV pulses while holding the potential at −60 mV. Cellular parameters were recorded initially and following each of the protocols run.

Three protocols were run on each cell: a voltage-clamp and two current-clamps, one run from the cell’s resting potential, and another run from holding the cell at −60 mV. The voltage-clamp protocol was performed from holding cells at a baseline of −60 mV. The current clamp protocol consisted of 20 pA current steps spaced between −100 to 480 pA, repeated 5 times. Neuronal recordings that were used for analysis were stable across the entire recording session, as measured by monitoring of consistent series resistance and resting potential. All spike analysis was performed in MATLAB vR2022b with custom scripts. These scripts are available in the following GitHub repository: https://github.com/DannyLasky/PatchClamp

### Electrophysiology Data Analysis

Resistance, capacitance, and resting potential were assessed initially and following each of the protocols run. As the data were consistent across the four time points, the metrics for each cell were averaged across time points. All spikes were detected from traces by an upward slope that crossed 30 mV/ms, followed in the next 10 ms by a downward slope that crossed −15 mV/ms. Each spike was additionally required to pass above 0 mV. Spontaneous and rebound spikes were omitted from analysis, which were collectively defined as spikes occurring outside of a depolarizing current application window.

Additional computations were performed when analyzing the first spike fired at rheobase, defined as the minimum current application necessary to elicit a spike. The maximum in the second derivative was found within a window spanning from the 30 mV/ms up cross, used for our spike detection, to 3 ms prior. This effectively located the greatest curvature of the spike, a marker of spike initiation. We defined the voltage of the original signal at this time point as “threshold,” the voltage at which the cell fired action potentials. The end of the spike was determined by when the spike crossed back under threshold. Latency was defined as the time from the beginning of current application to the spike threshold. We computed amplitude as the voltage from the threshold to the peak of the spike. Finally, we define width as the full width at half amplitude. On rare occasions (<1% of data), a first spike fired at rheobase did not cross back under its threshold. In this case, width could not be computed, so the spike was excluded from the additional rheobase calculations.

Voltage-clamp traces were leak subtracted. The smallest negative step of −10 mV was used as the best measure of passive current flowing through leak channels and capacitance. The current at this voltage step was scaled in a linear ohmic manner dependent on the voltage applied and then subtracted from the original signal. Following leak subtraction, the traces were zeroed using the current prior to applying a voltage step. Maximum inward and outward current were computed from the first 10 ms following voltage application. Steady-state currents for the IV curves were computed from the average current in the last third of the voltage application. Importantly, these voltage-clamp data were not space-clamped and do not properly account for the arborization of neurons. No liquid junction offsets were applied.

### Statistics

Datasets involving two groups were analyzed by unpaired Student’s t-test. For datasets with three groups, Brown-Forsythe and Welch one-way ANOVA was used. For datasets with four groups, two-way ANOVA with Tukey’s Multiple comparisons test was used. Sholl analyses were statistically assessed by a two-way ANOVA of the area under the curves.

Data for intrinsic excitability (Fig.4G–I, Fig. S5) and rheobase parameters (Fig.5, Fig. S7) were non-normal and analyzed appropriately with Kruskal–Wallis and post-hoc Dunn’s tests for the tau data and Mann Whitney U tests for the amyloid data. Data for spike count across current step, maximum inward and outward currents, and IV curves were analyzed with repeated measures two-way ANOVA with Geisser–Greenhouse correction^71^ and post-hoc Tukey tests. One or more spike and exactly one spike data were analyzed via F-Test, fitting one-phase association curves through the origin to one plus spike plots and fitting lines through the origin to exactly one spike plots. For all statistics, p < 0.05 was defined as significant. Asterisks on figures denote the following significance levels: * p<0.05, ** p<0.01, *** p<0.001, **** p<0.0001.

## Supplemental Figures

**Figure S1:** A) Schematic of neuron culture timeline. Neurons were isolated from postnatal day 0 mice. Neurons were seeded with 3 μg/mL PFF at day *in vitro* (DIV) 7. Cultures were maintained until at least DIV 25 before experimentation. B) Immunofluorescence of phosphorylated tau (AT-8) in Tau neurons seeded with PFF seeded or PBS control (n=25–28 cells) (scale bar 100 μm). Data shown as mean ± SEM. Significance determined by Student’s t-test with Welch’s correction. C) Schematic of AdipoRon treatment times used in this paper. Protein phosphorylation events were measured after 10-minute treatment. Mitochondrial membrane potential was assessed 2 hours after treatment. Tau clearance, mitochondrial morphology, neuronal architecture, and electrophysiology were measured after 24-hour treatment.

**Figure S2:** Representative immunofluorescence images of total protein levels of AMPK and GSK3b. Quantification is shown in Fig 1.

**Fig S3:** A) Volcano plots displaying the differentially expressed genes in RNAseq data. Comparisons shown are Tau vs. WT neurons, Tau neurons AR vs. Control, and WT neurons AR vs. Control. B) Venn diagram showing overlap between differentially expressed genes in the 3 comparisons shown in A. C) Venn diagram showing the overlap between enriched KEGG pathways in the 3 comparisons shown in A. D) Complete STRING diagram from the Neuroactive Ligand Receptor interaction pathway. This diagram was curated to show only the large networks in Fig.2C.

**Fig S4:** Quantification of lipid droplet number from neurons shown in Fig.3I. Data shown as mean ± SEM. Heatmaps of Pyrimidine metabolism and amino/ nucleotide sugar metabolism GSEA pathways in AR-treated WT and Tau neurons.

**Fig S5.** A) Representative immunofluorescence images of total protein levels of JNK. Quantification shown in Fig.4D. B–D) Resistance (B), capacitance (C), and resting potential (D) for APP/PS1 and WT neurons. Box plots display the median, interquartile range, and whiskers extending to values within the interquartile range multiplied by 1.5. Statistics were performed using Mann–Whitney U tests. D) Spike counts measured in response to stepwise current applications of 20 pA from −100–480 pA for each cell from rest (left) and from −60 mV (right). Statistics were performed using repeated measures two-way ANOVA with Geisser–Greenhouse correction and post-hoc Tukey tests. Error bars are SEM.

**Fig S6:** The percentage of cells firing one or more spikes (A) or exactly one spike (B) was measured across current steps. Lines were compared in a pairwise manner using F-tests of a one-phase association curve passing through the origin for one or more spike plots, and a line through the origin for the exactly one spike plots. Bonferroni corrections were applied to account for the multiple pairwise comparisons. Significance between groups is displayed as paired colored blocks. Sample sizes are: WT: 64, WT+AR: 13, Tau: 40, Tau+AR: 31. C) Action potential spike counts for APP/PS1 and WT neurons were measured in response to 30 stepwise current applications of 20 pA from −100–480 pA for each cell from rest (left) and from −60 mV (right). Statistics were performed using repeated measures two-way ANOVA with Geisser–Greenhouse correction and post-hoc Tukey tests. D-E) The percentage of cells firing one or more spikes (D) or exactly one spike (E) was measured across current steps. Lines were compared in a pairwise manner using F-tests of a one-phase association curve passing through the origin for one or more spike plots, and a line through the origin for the exactly one spike plots. Significance between groups is displayed as paired colored blocks. F-H) Spike latency (F), amplitude (G), and width (H) for the first spike fired at rheobase were collected from current-clamp protocols performed for each cell from rest (left) and from −60 mV (right). Spike width was measured from the full width at half amplitude. Box plots display the median, interquartile range, and whiskers extending to values within the interquartile range multiplied by 1.5. Statistics were performed using Mann–Whitney U tests. Error bars are SEM. Sample sizes for A–F and J–L are WT: 42, Amy: 28. Sample sizes for G–I are WT: 28, Amy: 17 from rest and WT: 35, Amy: 20 from −60 mV. (* p<0.05, *** p<0.001, **** p<0.0001).

**Fig. S7:** The maximum inward current, outward current,and IV curves in APP/PS1 and WT neurons in response to stepwise voltage injections of 10 mV from 0–100 mV. Statistics were performed using repeated measures two-way ANOVA with Geisser–Greenhouse correction and post-hoc Tukey tests. Error bars are SEM. Sample sizes for A–F and J–L are WT: 42, Amy: 28. Sample sizes for G–I are WT: 28, Amy: 17 from rest and WT: 35, Amy: 20 from −60 mV.

## References

1. Nelson PT, Alafuzoff I, Bigio EH, et al. Correlation of Alzheimer Disease Neuropathologic Changes With Cognitive Status: A Review of the Literature. J Neuropathol Exp Neurol. 2012;71(5):362–381. doi:10.1097/NEN.0b013e31825018f7

2. Tanner JA, Rabinovici GD. Relationship Between Tau and Cognition in the Evolution of Alzheimer’s Disease: New Insights from Tau PET. J Nucl Med. 2021;62(5):612–613. doi:10.2967/jnumed.120.257824

3. Xia Y, Prokop S, Giasson BI. “Don’t Phos Over Tau”: recent developments in clinical biomarkers and therapies targeting tau phosphorylation in Alzheimer’s disease and other tauopathies. Mol Neurodegeneration. 2021;16(1):37. doi:10.1186/s13024-021-00460-5

4. Orr ME, Sullivan AC, Frost B. A Brief Overview of Tauopathy: Causes, Consequences, and Therapeutic Strategies. Trends in Pharmacological Sciences. 2017;38(7):637–648. doi:10.1016/j.tips.2017.03.011

5. Toledo JB, Arnold M, Kastenmuller G, et al. Metabolic network failures in Alzheimer’s disease: A biochemical road map. Alzheimers Dement. 2017;13(9):965–984. doi:10.1016/j.jalz.2017.01.020

6. Souder DC, Dreischmeier IA, Smith AB, et al. Rhesus monkeys as a translational model for late-onset Alzheimer’s disease. Aging Cell. 2021;20(6). doi:10.1111/acel.13374

7. Perez MJ, Jara C, Quintanilla RA. Contribution of Tau Pathology to Mitochondrial Impairment in Neurodegeneration. Front Neurosci. 2018;12:441. doi:10.3389/fnins.2018.00441

8. Camandola S, Mattson MP. Brain metabolism in health, aging, and neurodegeneration. EMBO J. 2017;36(11):1474–1492. doi:10.15252/embj.201695810

9. Cunnane SC, Trushina E, Morland C, et al. Brain energy rescue: an emerging therapeutic concept for neurodegenerative disorders of ageing. Nat Rev Drug Discov. 2020;19(9):609–633. doi:10.1038/s41573-020-0072-x

10. Funcke JB, Scherer PE. Beyond adiponectin and leptin: adipose tissue-derived mediators of inter-organ communication. J Lipid Res. 2019;60(10):1648–1684. doi:10.1194/jlr.R094060

11. Wang ZV, Scherer PE. Adiponectin, the past two decades. J Mol Cell Biol. 2016;8(2):93–100. doi:10.1093/jmcb/mjw011

12. Li X, Zhang D, Vatner DF, et al. Mechanisms by which adiponectin reverses high fat diet-induced insulin resistance in mice. Proc Natl Acad Sci U S A. 2020;117(51):32584–32593. doi:10.1073/pnas.1922169117

13. Li N, Zhao S, Zhang Z, et al. Adiponectin preserves metabolic fitness during aging. Elife. 2021;10:e65108. doi:10.7554/eLife.65108

14. Ding Q, Ash C, Mracek T, Merry B, Bing C. Caloric restriction increases adiponectin expression by adipose tissue and prevents the inhibitory effect of insulin on circulating adiponectin in rats. J Nutr Biochem. 2012;23(8):867–874. doi:10.1016/j.jnutbio.2011.04.011

15. Miller KN, Burhans MS, Clark JP, et al. Aging and caloric restriction impact adipose tissue, adiponectin, and circulating lipids. Aging Cell. 2017;16(3):497–507. doi:10.1111/acel.12575

16. Kita S, Fukuda S, Maeda N, Shimomura I. Native adiponectin in serum binds to mammalian cells expressing T-cadherin, but not AdipoRs or calreticulin. eLife. 2019;8:e48675. doi:10.7554/eLife.48675

17. Thundyil J, Pavlovski D, Sobey CG, Arumugam TV. Adiponectin receptor signalling in the brain. Br J Pharmacol. 2012;165(2):313–327. doi:10.1111/j.1476-5381.2011.01560.x

18. Formolo DA, Cheng T, Yu J, Kranz GS, Yau SY. Central Adiponectin Signaling - A Metabolic Regulator in Support of Brain Plasticity. Brain Plast. 2022;8(1):79–96. doi:10.3233/BPL-220138

19. Lee TH, Ahadullah, Christie BR, et al. Chronic AdipoRon Treatment Mimics the Effects of Physical Exercise on Restoring Hippocampal Neuroplasticity in Diabetic Mice. Mol Neurobiol. 2021;58(9):4666–4681. doi:10.1007/s12035-021-02441-7

20. Liu Y, Palanivel R, Rai E, et al. Adiponectin Stimulates Autophagy and Reduces Oxidative Stress to Enhance Insulin Sensitivity During High-Fat Diet Feeding in Mice. Diabetes. 2015;64(1):36–48. doi:10.2337/db14-0267

21. Okada-Iwabu M, Yamauchi T, Iwabu M, et al. A small-molecule AdipoR agonist for type 2 diabetes and short life in obesity. Nature. 2013;503(7477):493–499. doi:10.1038/nature12656

22. He K, Nie L, Ali T, et al. Adiponectin alleviated Alzheimer-like pathologies via autophagy-lysosomal activation. Aging Cell. 2021;20(12). doi:10.1111/acel.13514

23. Wang C, Chang Y, Zhu J, et al. AdipoRon mitigates tau pathology and restores mitochondrial dynamics via AMPK-related pathway in a mouse model of Alzheimer’s disease. Experimental Neurology. 2023;363:114355. doi:10.1016/j.expneurol.2023.114355

24. Guo JL, Lee VMY. Seeding of Normal Tau by Pathological Tau Conformers Drives Pathogenesis of Alzheimer-like Tangles. Journal of Biological Chemistry. 2011;286(17):15317–15331. doi:10.1074/jbc.M110.209296

25. Jicha GA, Bowser R, Kazam IG, Davies P. Alz-50 and MC-1, a new monoclonal antibody raised to paired helical filaments, recognize conformational epitopes on recombinant tau. J Neurosci Res. 1997;48(2):128–132. doi:10.1002/(SICI)1097-4547(19970415)48:2<lt;128::AID-JNR5>3.0.CO;2-E

26. Zhou Q, Li S, Li M, et al. Human tau accumulation promotes glycogen synthase kinase-3β acetylation and thus upregulates the kinase: A vicious cycle in Alzheimer neurodegeneration. eBioMedicine. 2022;78:103970. doi:10.1016/j.ebiom.2022.103970

27. Wang M, Jo J, Song J. Adiponectin improves long-term potentiation in the 5XFAD mouse brain. Sci Rep. 2019;9(1):8918. doi:10.1038/s41598-019-45509-0

28. Wang Y, Lam JB, Lam KSL, et al. Adiponectin Modulates the Glycogen Synthase Kinase-3β/β-Catenin Signaling Pathway and Attenuates Mammary Tumorigenesis of MDA-MB-231 Cells in Nude Mice. Cancer Research. 2006;66(23):11462–11470. doi:10.1158/0008-5472.CAN-06-1969

29. Horike N, Sakoda H, Kushiyama A, et al. AMP-activated Protein Kinase Activation Increases Phosphorylation of Glycogen Synthase Kinase 3β and Thereby Reduces cAMP-responsive Element Transcriptional Activity and Phosphoenolpyruvate Carboxykinase C Gene Expression in the Liver. Journal of Biological Chemistry. 2008;283(49):33902–33910. doi:10.1074/jbc.M802537200

30. Park SY, Lee YK, Lee WS, Park OJ, Kim YM. The involvement of AMPK/GSK3-beta signals in the control of metastasis and proliferation in hepato-carcinoma cells treated with anthocyanins extracted from Korea wild berry Meoru. BMC Complement Altern Med. 2014;14(1):109. doi:10.1186/1472-6882-14-109

31. Miller KN, Clark JP, Martin SA, et al. PGC-1a integrates a metabolism and growth network linked to caloric restriction. Aging Cell. 2019;18(5). doi:10.1111/acel.12999

32. Koutsodendris N, Blumenfeld J, Agrawal A, et al. Neuronal APOE4 removal protects against tau-mediated gliosis, neurodegeneration and myelin deficits. Nat Aging. 2023;3(3):275–296. doi:10.1038/s43587-023-00368-3

33. Ke YD, Chan G, Stefanoska K, et al. CNS cell type-specific gene profiling of P301S tau transgenic mice identifies genes dysregulated by progressive tau accumulation. J Biol Chem. 2019;294(38):14149–14162. doi:10.1074/jbc.RA118.005263

34. Miao Q, Ge M, Huang L. Up-regulation of GBP2 is Associated with Neuronal Apoptosis in Rat Brain Cortex Following Traumatic Brain Injury. Neurochem Res. 2017;42(5):1515–1523. doi:10.1007/s11064-017-2208-x

35. Dinman JD. Pathways to Specialized Ribosomes: The Brussels Lecture. Journal of Molecular Biology. 2016;428(10, Part B):2186–2194. doi:10.1016/j.jmb.2015.12.021

36. Ferretti MB, Karbstein K. Does functional specialization of ribosomes really exist? RNA. 2019;25(5):521–538. doi:10.1261/rna.069823.118

37. Wong TP, Howland JG, Wang YT. NMDA Receptors and Disease+C464. In: Squire LR, ed. Encyclopedia of Neuroscience. Academic Press; 2009:1177–1182. doi:10.1016/B978-008045046-9.01223-7

38. Nitsch RM, Deng M, Growdon JH, Wurtman RJ. Serotonin 5-HT2a and 5-HT2c receptors stimulate amyloid precursor protein ectodomain secretion. J Biol Chem. 1996;271(8):4188–4194. doi:10.1074/jbc.271.8.4188

39. Jahreis K, Brüge A, Borsdorf S, et al. Amisulpride as a potential disease-modifying drug in the treatment of tauopathies. Alzheimer’s & Dementia. 2023;19(12):5482–5497. doi:10.1002/alz.13090

40. Yung YC, Stoddard NC, Mirendil H, Chun J. Lysophosphatidic acid (LPA) signaling in the nervous system. Neuron. 2015;85(4):669–682. doi:10.1016/j.neuron.2015.01.009

41. Hackelberg S, Oliver D. Metabotropic Acetylcholine and Glutamate Receptors Mediate PI(4,5)P2 Depletion and Oscillations in Hippocampal CA1 Pyramidal Neurons in situ. Sci Rep. 2018;8(1):12987. doi:10.1038/s41598-018-31322-8

42. Yim WWY, Mizushima N. Lysosome biology in autophagy. Cell Discov. 2020;6(1):6. doi:10.1038/s41421-020-0141-7

43. Hamano T, Enomoto S, Shirafuji N, et al. Autophagy and Tau Protein. IJMS. 2021;22(14):7475. doi:10.3390/ijms22147475

44. Uddin MdS, Stachowiak A, Mamun AA, et al. Autophagy and Alzheimer’s Disease: From Molecular Mechanisms to Therapeutic Implications. Front Aging Neurosci. 2018;10:04. doi:10.3389/fnagi.2018.00004

45. Lee JH, Yang DS, Goulbourne CN, et al. Faulty autolysosome acidification in Alzheimer’s disease mouse models induces autophagic build-up of Aβ in neurons, yielding senile plaques. Nat Neurosci. 2022;25(6):688–701. doi:10.1038/s41593-022-01084-8

46. Klionsky DJ, Abdelmohsen K, Abe A, et al. Guidelines for the use and interpretation of assays for monitoring autophagy (3rd edition). Autophagy. 2016;12(1):1–222. doi:10.1080/15548627.2015.1100356

47. Jang M, Park R, Kim H, et al. AMPK contributes to autophagosome maturation and lysosomal fusion. Sci Rep. 2018;8(1):12637. doi:10.1038/s41598-018-30977-7

48. Yim WWY, Mizushima N. Lysosome biology in autophagy. Cell Discov. 2020;6(1):6. doi:10.1038/s41421-020-0141-7

49. Tilija Pun N, Park PH. Role of p62 in the suppression of inflammatory cytokine production by adiponectin in macrophages: Involvement of autophagy and p21/Nrf2 axis. Sci Rep. 2017;7(1):393. doi:10.1038/s41598-017-00456-6

50. Onyango IG, Dennis J, Khan SM. Mitochondrial Dysfunction in Alzheimer’s Disease and the Rationale for Bioenergetics Based Therapies. Aging and disease. 2016;7(2):201. doi:10.14336/AD.2015.1007

51. Johnson ECB, Dammer EB, Duong DM, et al. Large-scale proteomic analysis of Alzheimer’s disease brain and cerebrospinal fluid reveals early changes in energy metabolism associated with microglia and astrocyte activation. Nat Med. 2020;26(5):769–780. doi:10.1038/s41591-020-0815-6

52. Lozoya OA, Xu F, Grenet D, et al. A brain-specific *pgc1* α fusion transcript affects gene expression and behavioural outcomes in mice. Life Sci Alliance. 2021;4(12):e202101122. doi:10.26508/lsa.202101122

53. St-Pierre J, Drori S, Uldry M, et al. Suppression of Reactive Oxygen Species and Neurodegeneration by the PGC-1 Transcriptional Coactivators. Cell. 2006;127(2):397–408. doi:10.1016/j.cell.2006.09.024

54. Martínez-Redondo V, Pettersson AT, Ruas JL. The hitchhiker’s guide to PGC-1α isoform structure and biological functions. Diabetologia. 2015;58(9):1969–1977. doi:10.1007/s00125-015-3671-z

55. Monzel AS, Enríquez JA, Picard M. Multifaceted mitochondria: moving mitochondrial science beyond function and dysfunction. Nat Metab. 2023;5(4):546–562. doi:10.1038/s42255-023-00783-1

56. Farmer BC, Walsh AE, Kluemper JC, Johnson LA. Lipid Droplets in Neurodegenerative Disorders. Front Neurosci. 2020;14:742. doi:10.3389/fnins.2020.00742

57. Brandt R, Lee G. Functional organization of microtubule-associated protein tau. Identification of regions which affect microtubule growth, nucleation, and bundle formation in vitro. Journal of Biological Chemistry. 1993;268(5):3414–3419. doi:10.1016/S0021-9258(18)53710-8

58. Mijalkov M, Volpe G, Fernaud-Espinosa I, DeFelipe J, Pereira JB, Merino-Serrais P. Dendritic spines are lost in clusters in Alzheimer’s disease. Sci Rep. 2021;11(1):12350. doi:10.1038/s41598-021-91726-x

59. D’Andrea MR, Ilyin S, Plata-Salaman CR. Abnormal patterns of microtubule-associated protein-2 (MAP-2) immunolabeling in neuronal nuclei and Lewy bodies in Parkinson’s disease substantia nigra brain tissues. Neuroscience Letters. 2001;306(3):137–140. doi:10.1016/S0304-3940(01)01811-0

60. Chang L, Jones Y, Ellisman MH, Goldstein LSB, Karin M. JNK1 Is Required for Maintenance of Neuronal Microtubules and Controls Phosphorylation of Microtubule-Associated Proteins. Developmental Cell. 2003;4(4):521–533. doi:10.1016/S1534-5807(03)00094-7

61. Oliva AA, Atkins CM, Copenagle L, Banker GA. Activated c-Jun N-Terminal Kinase Is Required for Axon Formation. J Neurosci. 2006;26(37):9462–9470. doi:10.1523/JNEUROSCI.2625-06.2006

62. Atkinson PJ, Cho CH, Hansen MR, Green SH. Activity of all JNK isoforms contributes to neurite growth in spiral ganglion neurons. Hearing Research. 2011;278(1-2):77–85. doi:10.1016/j.heares.2011.04.011

63. Miyazaki T, Bub JD, Uzuki M, Iwamoto Y. Adiponectin activates c-Jun NH2-terminal kinase and inhibits signal transducer and activator of transcription 3. Biochemical and Biophysical Research Communications. 2005;333(1):79–87. doi:10.1016/j.bbrc.2005.05.076

64. Sun Q, Jiang YQ, Lu MC. Topographic heterogeneity of intrinsic excitability in mouse hippocampal CA3 pyramidal neurons. Journal of Neurophysiology. 2020;124(4):1270–1284. doi:10.1152/jn.00147.2020

65. Hampel H, Hardy J, Blennow K, et al. The Amyloid-β Pathway in Alzheimer’s Disease. Mol Psychiatry. 2021;26(10):5481–5503. doi:10.1038/s41380-021-01249-0

66. Choi SR, Lim JH, Kim MY, et al. Adiponectin receptor agonist AdipoRon decreased ceramide, and lipotoxicity, and ameliorated diabetic nephropathy. Metabolism: clinical and experimental. 2018;85:348–360. doi:10.1016/j.metabol.2018.02.004

67. Cheng A, Wan R, Yang JL, et al. Involvement of PGC-1alpha in the formation and maintenance of neuronal dendritic spines. Nat Commun. 2012;3:1250. doi:10.1038/ncomms2238

68. Yang HQ, Echeverry FA, ElSheikh A, et al. Subcellular trafficking and endocytic recycling of KATP channels. Am J Physiol Cell Physiol. 2022;322(6):C1230–C1247. doi:10.1152/ajpcell.00099.2022

69. Khandelwal M, Manglani K, Upadhyay P, Azad M, Gupta S. AdipoRon induces AMPK activation and ameliorates Alzheimer’s like pathologies and associated cognitive impairment in APP/PS1 mice. Neurobiology of Disease. 2022;174:105876. doi:10.1016/j.nbd.2022.105876

70. Takeuchi H, Iba M, Inoue H, et al. P301S Mutant Human Tau Transgenic Mice Manifest Early Symptoms of Human Tauopathies with Dementia and Altered Sensorimotor Gating. PLoS One. 2011;6(6):e21050. doi:10.1371/journal.pone.0021050

71. Corsi A, Bombieri C, Valenti MT, Romanelli MG. Tau Isoforms: Gaining Insight into MAPT Alternative Splicing. Int J Mol Sci. 2022;23(23):15383. doi:10.3390/ijms232315383

