## Supplementary figures and images for "Reversal of neuronal tau pathology, metabolic dysfunction, and electrophysiological defects via adiponectin pathway-dependent AMPK activation"

### Supplemental Figure 1

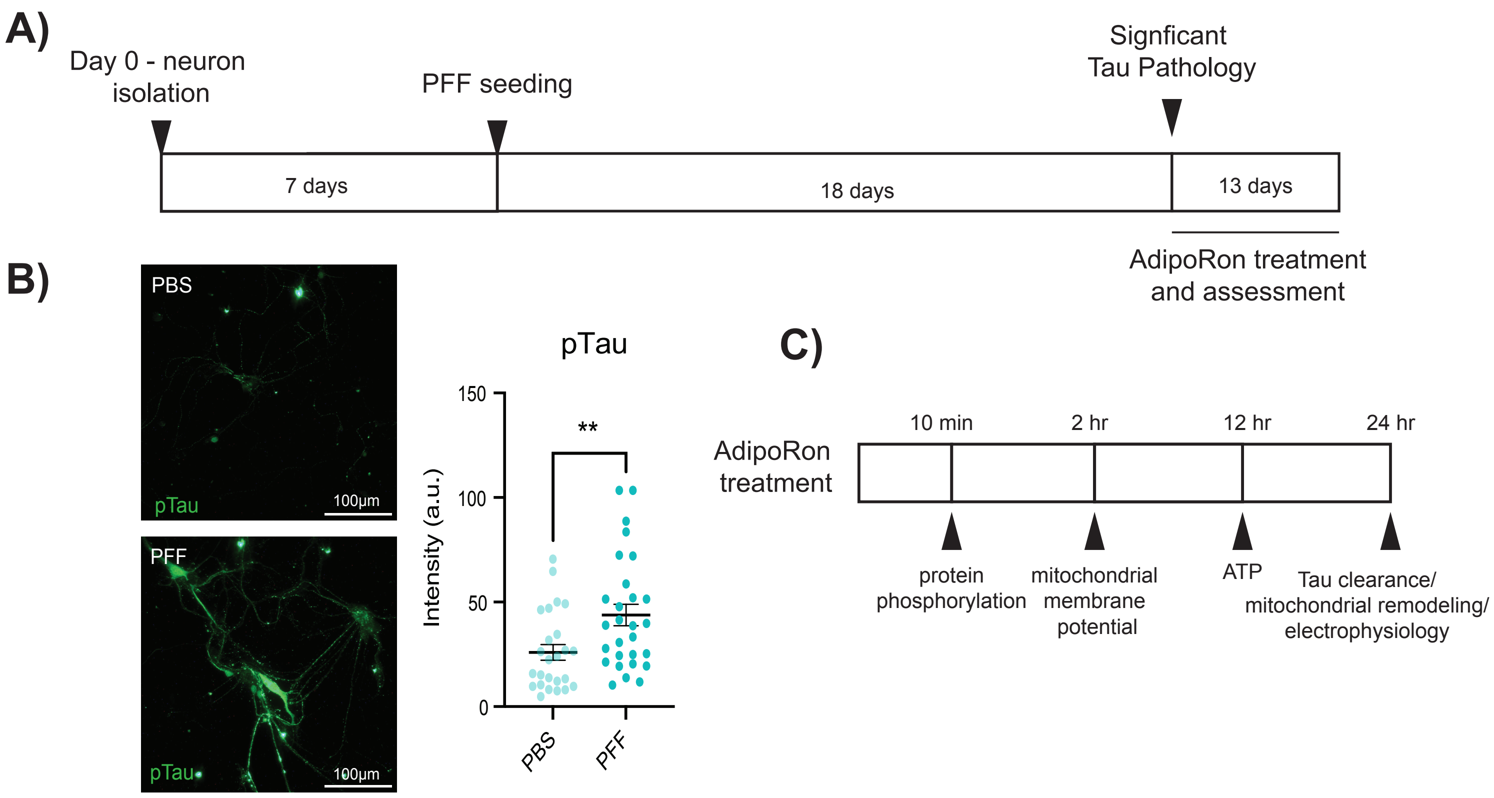

### Supplemental Figure 2

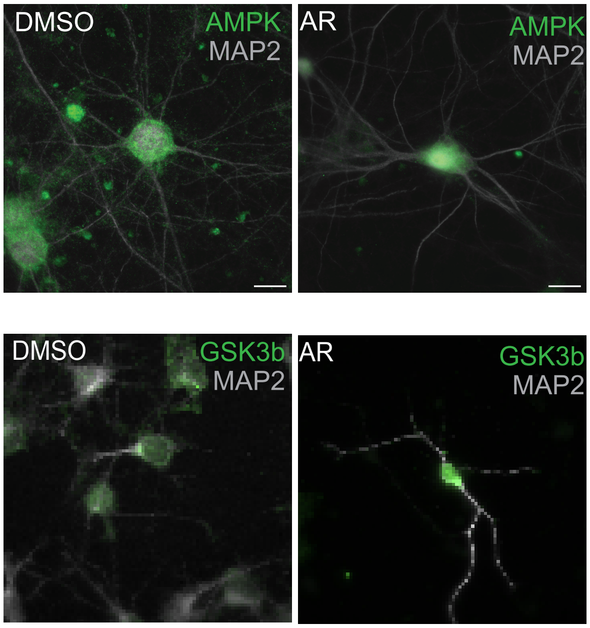

### Supplemental Figure 3

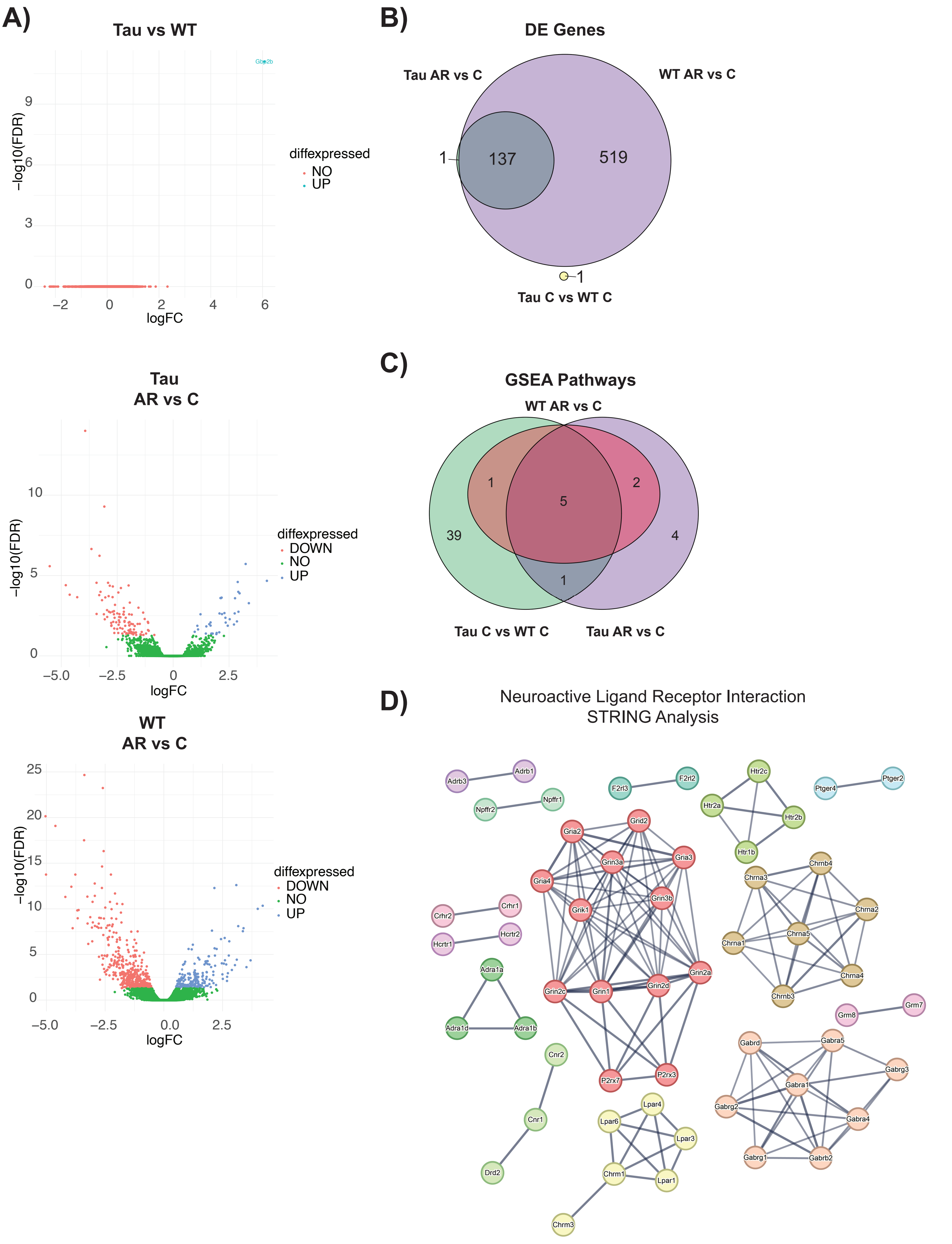

### Supplemental Figure 4

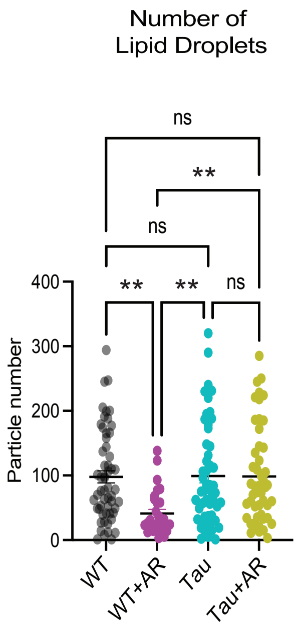

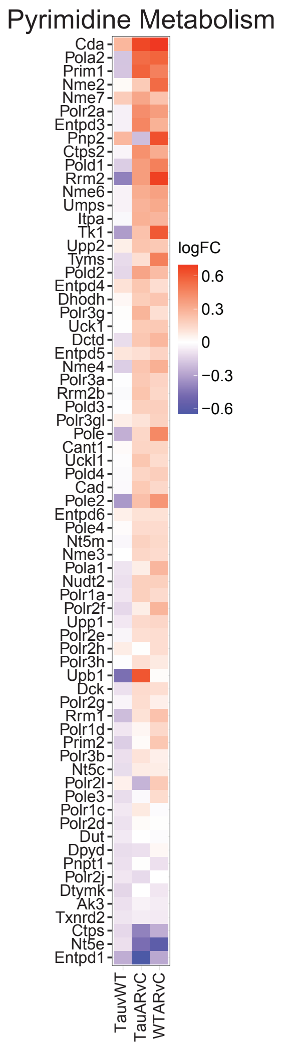

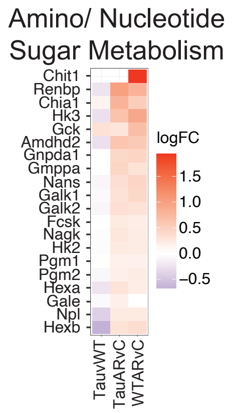

### Supplemental Figure 5

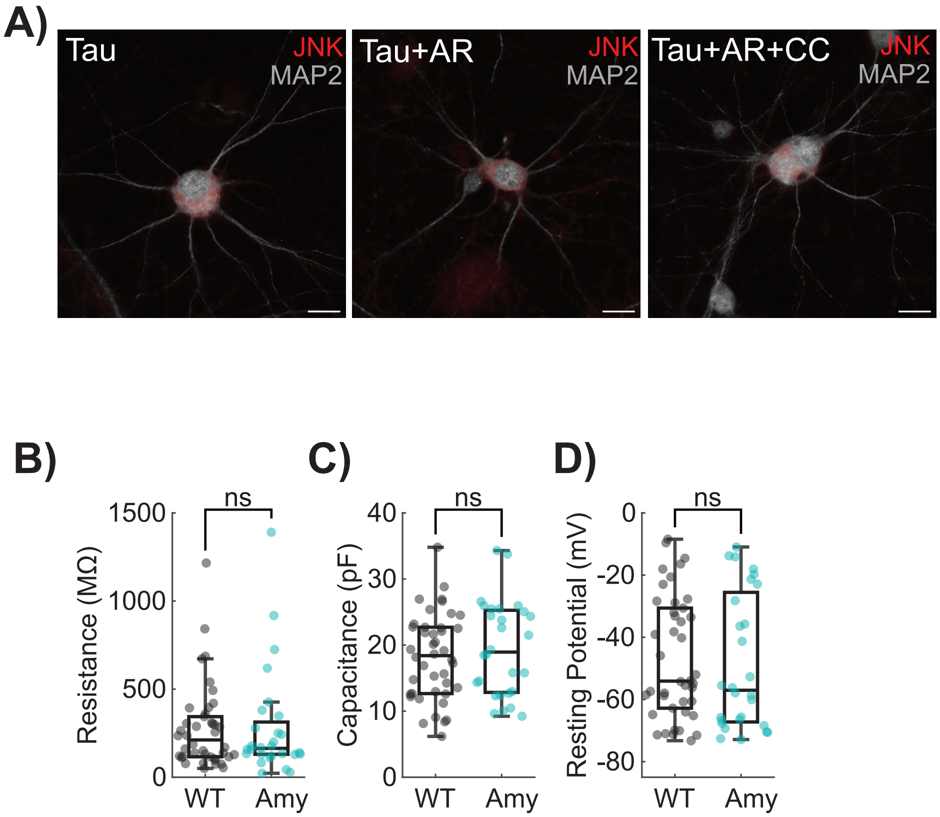

### Supplemental Figure 6

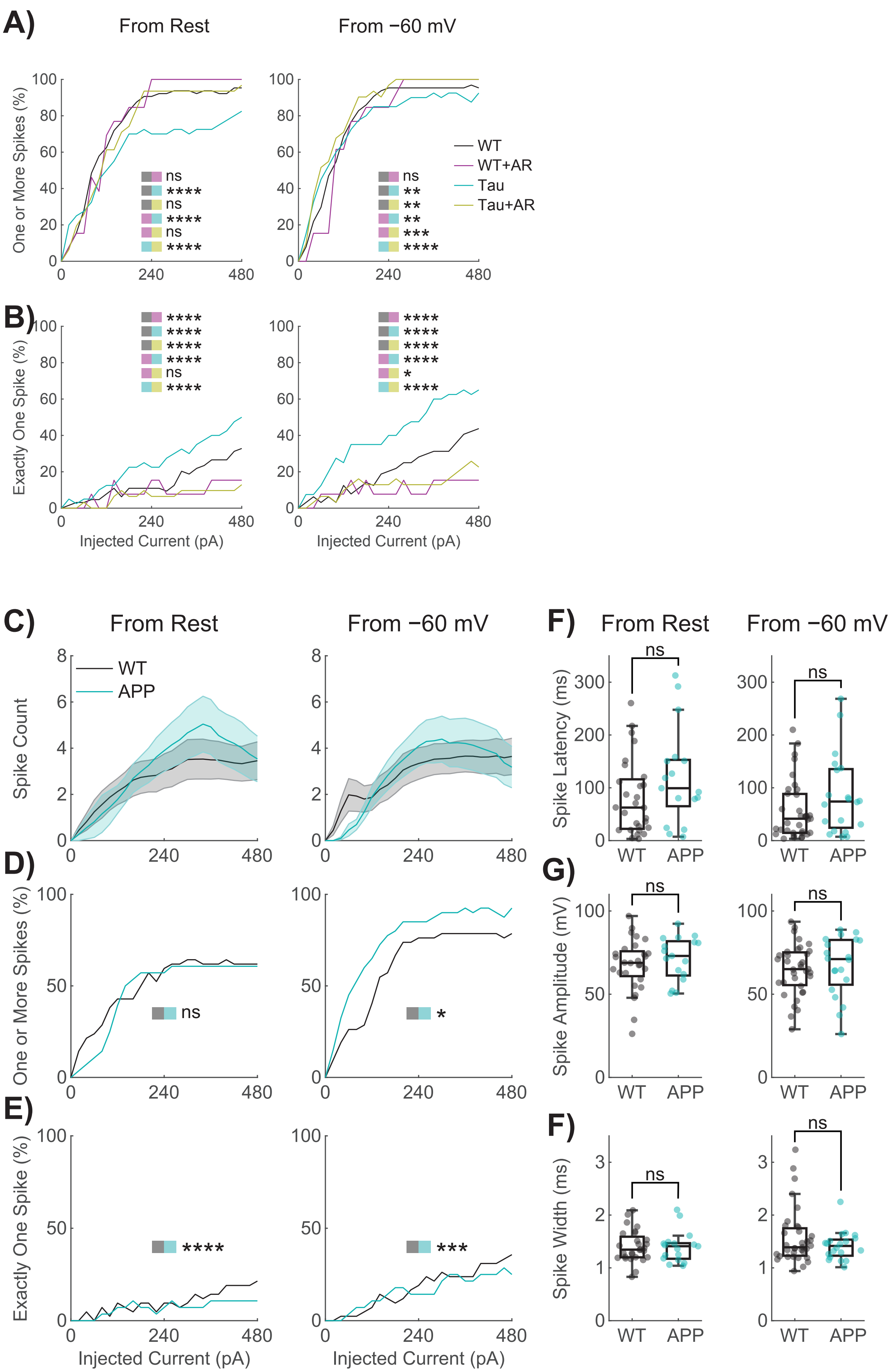

### Supplemental Figure 7

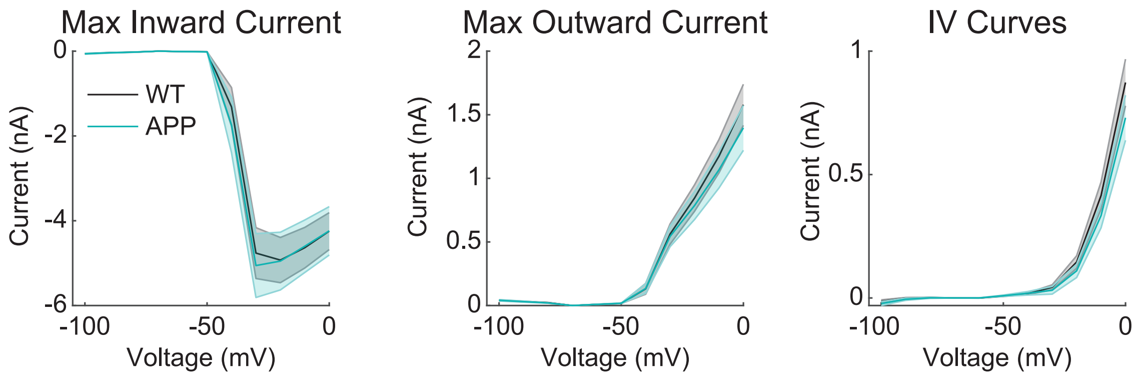
