## Supplemental Table 2 for "Reversal of neuronal tau pathology, metabolic dysfunction, and electrophysiological defects via adiponectin pathway-dependent AMPK activation"

| Voltage clamp measurement | Main group effect | Main Voltage effect | Group x Voltage interaction effect |
| --- | --- | --- | --- |
| Maximum Inward Current | F_(3,144)_=6.227, p=0.0005 | F_(1.827,263.1)_=298.8, p<0.0001 | F_(30,1440)_=4.709, p<0.0001 |
| Maximum Outward Current | F_(3,144)_=8.721, p<0.0001 | F_(1.174,169)_=450.3, p<0.0001 | F_(30,1440)_=7.820, p<0.0001 |
| IV Curve | F_(3,144)_ = 7.269, p=0.0001 | F_(1.132,163)_ = 258.2, p<0.0001 | F_(30,1440)_ = 6.175, p<0.0001 |

Table S2. AdipoRon changes voltage-gated channel activity in Tau neurons.
